# Increased neuron density in the midbrain of foveate birds results from profound change in tissue morphogenesis

**DOI:** 10.1101/2022.10.29.514341

**Authors:** Tania Rodrigues, Linda Dib, Émilie Bréthaut, Michel M. Matter, Lidia Matter-Sadzinski, Jean-Marc Matter

**Author notes:** Correspondance (T.R.) (J.-M.M.).

## Abstract

The increase of brain neuron number in relation with brain size is currently considered to be the major evolutionary path to high cognitive power in amniotes. However, how changes in neuron density did contribute to the evolution of the information-processing capacity of the brain remains unanswered. High neuron densities are seen as the main reason why the fovea located at the optical center of the retina is responsible for sharp vision in birds and primates. The emergence of foveal vision is considered as a breakthrough innovation in visual system evolution. We found that neuron densities in the largest visual center of the midbrain – i.e., the optic tectum – are two to four times higher in modern birds with one or two foveae compared to birds deprived of this specialty. Interspecies comparisons enabled us to identify elements of a hitherto unknown developmental process set up by foveate birds for increasing neuron density in the upper layers of their optic tectum. The late progenitor cells that generate these neurons proliferate in a ventricular zone that can expand only radially. In this particular context, the number of cells in ontogenetic columns increases, thereby setting the conditions for higher cell densities in the upper layers once neurons did migrate.

**Graphical Abstract:** 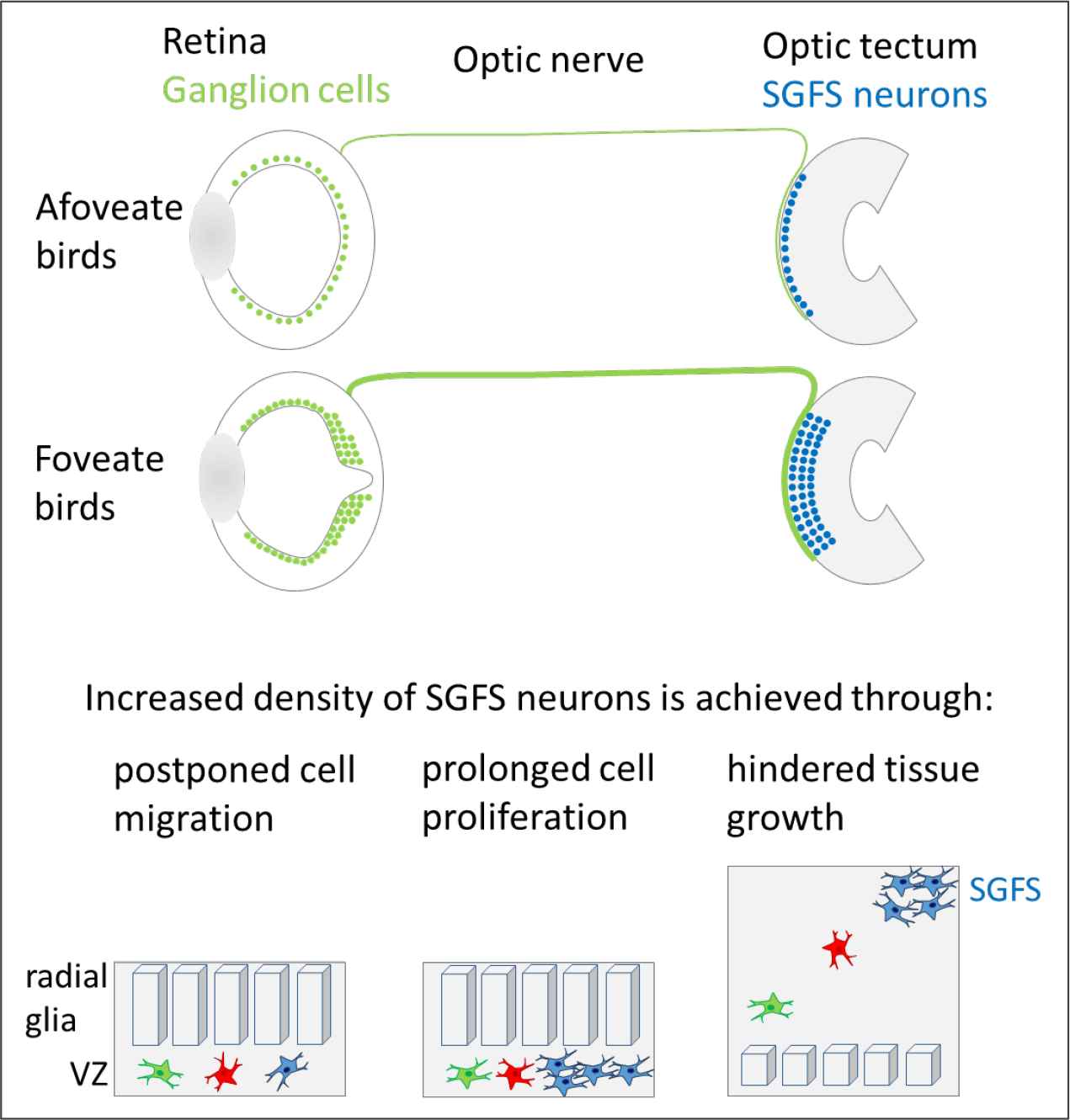

**Highlights:**

- The presence of a fovea is associated with increased neuron density in the optic tectum
- Fovea number and deepness influence neuron density in the optic tectum
- Increased neuron density in the upper layers of the optic tectum requires major developmental changes
- Postponed cell migration, prolonged cell proliferation, hindered tissue growth are necessary settings for achieving high neuron density

## INTRODUCTION

High visual acuity in primates is restricted to central vision, while acuity is low in the periphery. A major reason is that the density of retinal ganglion cells (RGCs) falls precipitously away from the fixation point, i.e., the fovea. About half of the ∼1.2×10^6^ RGCs in the human retina are located in the fovea, which represents less than 1% of the total retinal surface (Bringmann et al., 2018; Provis et al., 2013). RGCs whose bundled axons make up the optic nerve transmit visual information from the retina to the brain. While it is well known that the fovea makes primate visual perception unique among mammals, what is less well known is that this retinal specialization is of paramount importance for spatial and temporal acuities in birds of the Neoaves clade encompassing ∼95% of the ∼10’500 bird species (Bringmann, 2019; Jarvis et al., 2014; Prum et al., 2015). The recognition that the fovea is a region for acute vision dates back to the end of the 19th century (Ramón y Cajal, 1893; Slonaker, 1897). Despite this early interest, we know very little about neural signaling in the fovea.

In birds, the optic tectum (OT) is the largest visual center and its twin lobes form the roof of the midbrain. RGC axons terminate in the superficial layers of the contralateral lobe where they establish connections with upper neurons of the stratum griseum et fibrosum superficiale (SGFS). Retinal inputs are retinotopically distributed over the tectal surface. Electrophysiological recordings have shown that the foveal projection in the lateral geniculate nucleus, the superior colliculus and the primary visual cortex (V1) of primates and on the tectal surface of birds is a region of increased areal magnification (Clarke and Whitteridge, 1976; Cowey and Rolls, 1974; Daniel and Whitteridge, 1961; Malpeli and Baker, 1975; Rolls and Cowey, 1970; Rosa et al., 1997). It is assumed that because of the high number and high density of RGCs in their fovea, primates must have more neurons in V1 than non-primate mammals to account for the differences in acuity. Quantitative evaluations of cell number and density indicate that primates increase the number of neurons by increasing their density in the V1 area (Collins et al., 2010; Rockel et al., 1980; Srinivasan et al., 2015). However, the fact that in the Grey Mouse Lemur (*Microcebus murinus*) V1 occupies more than 20% of the neocortical area, the highest described in the primates to date (Ho et al., 2021) raises the possibility that increasing the V1 area could also contribute to increase the number of neurons. Thus, it is still unclear to what extent the increase of neuron density in the visual areas of the brain contributes to the enhanced representation of the fovea.

To address this question, we asked how birds adapted to an increased number of retinal inputs. First, we measured the surface density of neurons in the SGFS of birds who have one or two foveae or who are deprived of this retinal specialty. We found that cell densities were two to four times higher in the foveate birds. Although the retinal histology of less than one hundred bird species has been reported so far, we can reasonably assume that the fovea is one trait that distinguishes Neoaves from Galloanserae. Unlike the Galloanserae, which are precocial, the majority of modern birds are altricial, i.e., birds hatch with closed eyes and require prolonged parental care. Altriciality, by changes in the rate of development, may help to relax constraints on the duration of ontogenesis, a precondition for brain expansion and cognitive development. The Common Pigeon (*Columba livia domestica*) is altricial and the about three times higher density of RGCs in its retina compared to that in Chicken (*Gallus gallus domesticus*), primarily results from a prolonged expansion of the founder neural cell pool and a general delay of neurogenesis (Rodrigues et al., 2016). Our study uncovers that in the OT, in contrast to what happens in the retina, the onset of neurogenesis coincides in Common Pigeon and Chicken. Then, from the 7^th^ day of embryonic development (E7), there are major points of divergences between the two species. In Chicken, tectal cells stop dividing and newborn precursor cells rapidly migrate to establish the different outer layers (e.g., SGFS). In Common Pigeon, neurons differentiate while they get stuck in the ventricular zone (VZ) up to the end of the expansion of a late pool of progenitor cells committed to generate neural cells of the SGFS. Morphological, histological and transcriptomic comparisons of OT development in Common Pigeon and Chicken enabled us to identify elements of a hitherto unknown system deployed by modern birds for increasing neuron densities in the midbrain.

## RESULTS

The results are organized around two main findings about the adaptation of the optic tectum to foveal vision: (1) increased neuron density in the optic tectum of bird species with one or two foveae; (2) consistent with this morphological difference, the development of the optic tectum of foveate and afoveate birds differs dramatically.

### 1. Interspecies differences of cell densities in the optic tectum of Neoaves and Galloanserae

Behavioral tests show that birds have the best visual acuity when their retinas possess one or two foveae. The large majority of RGC axons that compose the optic nerve terminate in the SGFS of the OT where they establish connections with neurons in an orderly fashion according to a retinotopic mapping (Figure 1A). We asked whether there was a link between the number and architecture of fovea and the cell density in the SGFS. We selected twelve bird species with one or two foveae or without this retinal specialty. They were from seven taxonomic Orders with varying brain weight up to one order of magnitude (Table S1). Histological analysis of the foveae and cell counting in the OTs were performed on the same specimens. Although all the avian predators we examined have laterally placed eyes, except for the Tawny Owl (*Strix aluco*), they exhibit striking differences in terms of fovea organization (Figure 2). The Common Kestrel (*Falco tinnunculus*) has a central and a peripheral fovea, the Northern Goshawk (*Accipiter gentilis*), the Eurasian Sparrowhawk (*Accipiter nisus*) and the Black Kite (*Milvus migrans*) have only a central fovea and a peripheral area with, for some of them, high densities of RGCs and cones, whereas the Tawny Owl has no fovea. The retina of this truly nocturnal species is exclusively populated by rods and the density of RGCs is uniformly low (Figures 2, 3). Although, the Northern Goshawk, the Eurasian Sparrowhawk, the Grey Wagtail (*Motacilla cinerea*) and the Common Pigeon have different feeding habits and foraging behavior, their central fovea looks remarkably similar and none of these species have a peripheral fovea. The Common Swift (*Apus apus*) is among the fastest fliers, catching insects on the wing, yet, its single fovea is situated on the extreme periphery of the retina (Figure 2). To make things even more convoluted, the two swallows studied here, i.e., the Common House-Martin (*Delichon urbicum*) and the Barn Swallow (*Hirundo rustica*) have a central and a peripheral fovea, while their lifestyle and feeding habits are very similar to that of the swifts.

**FIGURE 1.**
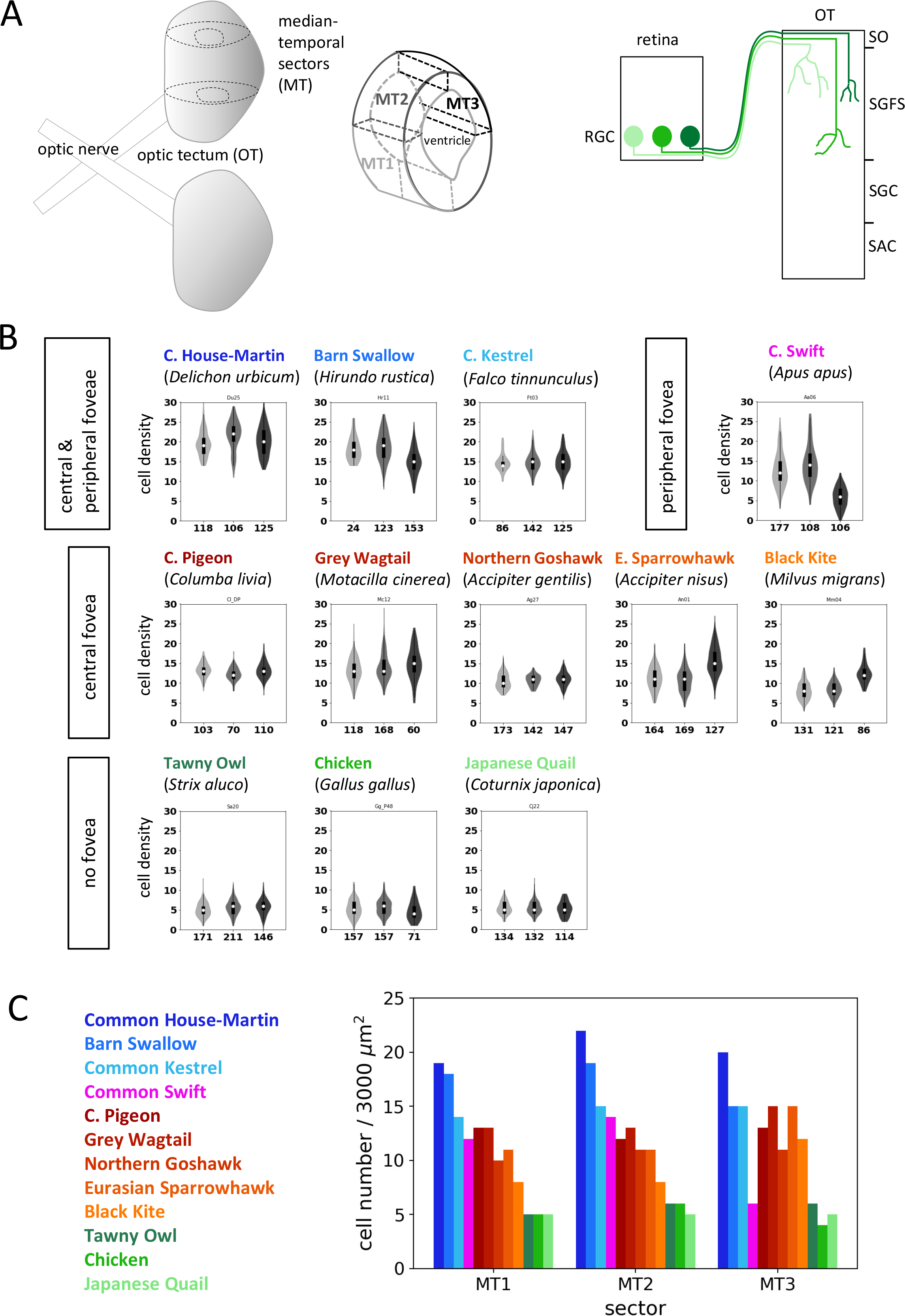
Cell density in the optic tectum. (A, left panel) the three median-temporal sectors of the optic tectum (OT) selected for cell counting. (A, right panel) Retinal ganglion cells (RGCs) innervate the superficial layers of the OT (SGFS) according to a retinotopic map. SO, stratum opticum, SGFS, stratum griseum et fibrosum superficiale, SGC, stratum griseum centrale, SAC, stratum album centrale. (B) Violin plots showing, for each studied species, the number of cells fitting in 50 μm x 60 μm frames (on the y-axis) used to screen the sectors MT1 (light gray), MT2 (middle gray) and MT3 (dark gray). The cell density is thus defined as the number of cells fitting in such a 3000 μm² counting frame. Values on the x-axis correspond to the number of counting frames used to screen each sector. There is one violin plot per species and per sector. Each violin plot shows the empirical distribution of the cell density obtained with a Gaussian kernel density estimation. Additionally, a more traditional box plot is overlaid: the white dot on each violin plot shows the median, the vertical black bar shows the interquartile range (IQR), whereas the black line shows the range between lower and upper whiskers. Lower (respectively upper) whisker is located at a 1.5 IQR distance measured down (respectively up) from the lower (respectively upper) quartile or at the smallest (respectively largest) data point. The values on the x-axis are the number of data points – i.e., the number of counting frames used to draw each violin plot as well as the overlying box plot. (C) Bar plot showing, for each studied species, the median cell densities (on the y-axis) across the three sectors of the median-temporal domain. The color of each bar is the color identifying the corresponding species.

**FIGURE 2.**
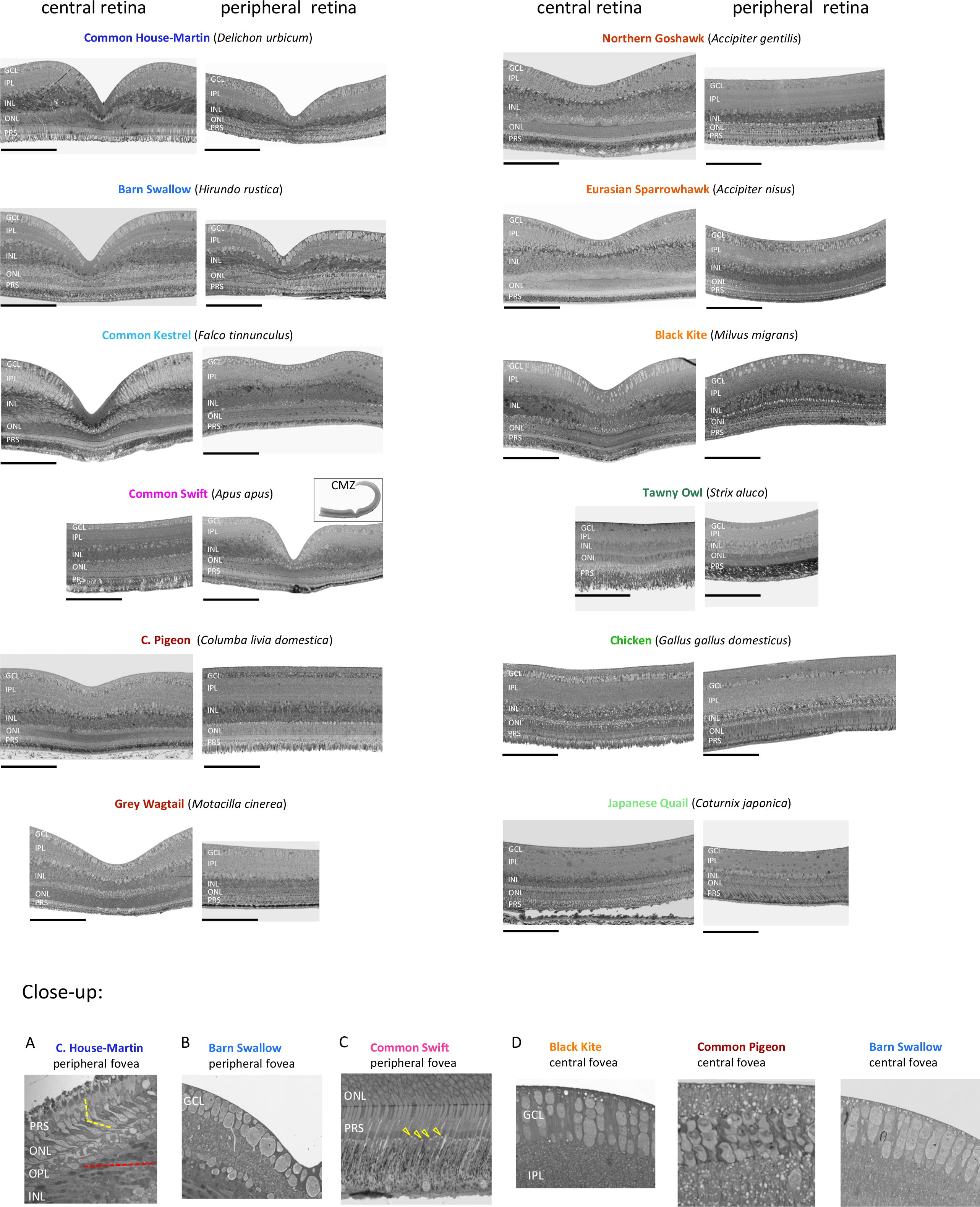
Fovea diversity in birds. Pictures of semi-thin cross-sections (1 μm thick) of the central and peripheral retinas of the twelve bird specimens investigated in this study (Table S1). All pictures (except the inset and the close-up pictures) are at the same scale. The foveate or afoveate peripheral retinal areas shown here are located in the temporal quadrant. Cell counts in the GCL and ONL are presented in the Figure 3. Of special note: (1) While the eye of the Common Kestrel is 2.2 times larger than that of the Barn Swallow (Table S1), the sizes, the deepness and the shape of the central foveae of these two species are identical. (2) In the Common Swift, the only fovea is located in the far periphery (inset), where the retina joins the ciliary marginal zone (CMZ). The pit is deep despite its eccentric position. (3) We did not find differences (e.g., deepness of the fovea, cell density in the GCL and ONL) between captive domestic pigeons and racing (homing) specimens, as well as between 10 weeks (P70) and 6 years old pigeons (our unpublished data). (4) While the eye of the male Northern Goshawk is 1.4 times larger than that of the male European Sparrowhawk (Table S1), the sizes, the deepness and the shapes of the central foveae of these two Accipiter species are identical. (5) The Tawny Owl is the only Neoaves of our collection with no fovea. The photoreceptors of this true nocturnal bird are rods with long outer segments (PRS). (A-D) Six close-up pictures illustrating special features of bird foveae. (A) The bending of the inner segments of cones lying within the pit (dashed yellow line). The inner segments tend to get aligned with rows of bipolar cells in the INL (dashed red line). (B) Somas of giant RGCs (surface areas ≥100µm^2^; Figure 3B) on the edge and within the pit of the peripheral fovea. (C) cone photoreceptors identifiable by the presence of an oil droplet in the outer segment (arrowheads). (D) Columnar organization of cells in the GCL is observed on the edges of both deep and shallow central foveae. It is, however, less frequent in peripheral foveae. PRS, outer segment of photoreceptor; ONL, outer nuclear layer; INL, inner nuclear layer; IPL, inner plexiform layer; GCL, ganglion cell layer. Scale bars: 200 µm.

**FIGURE 3.**
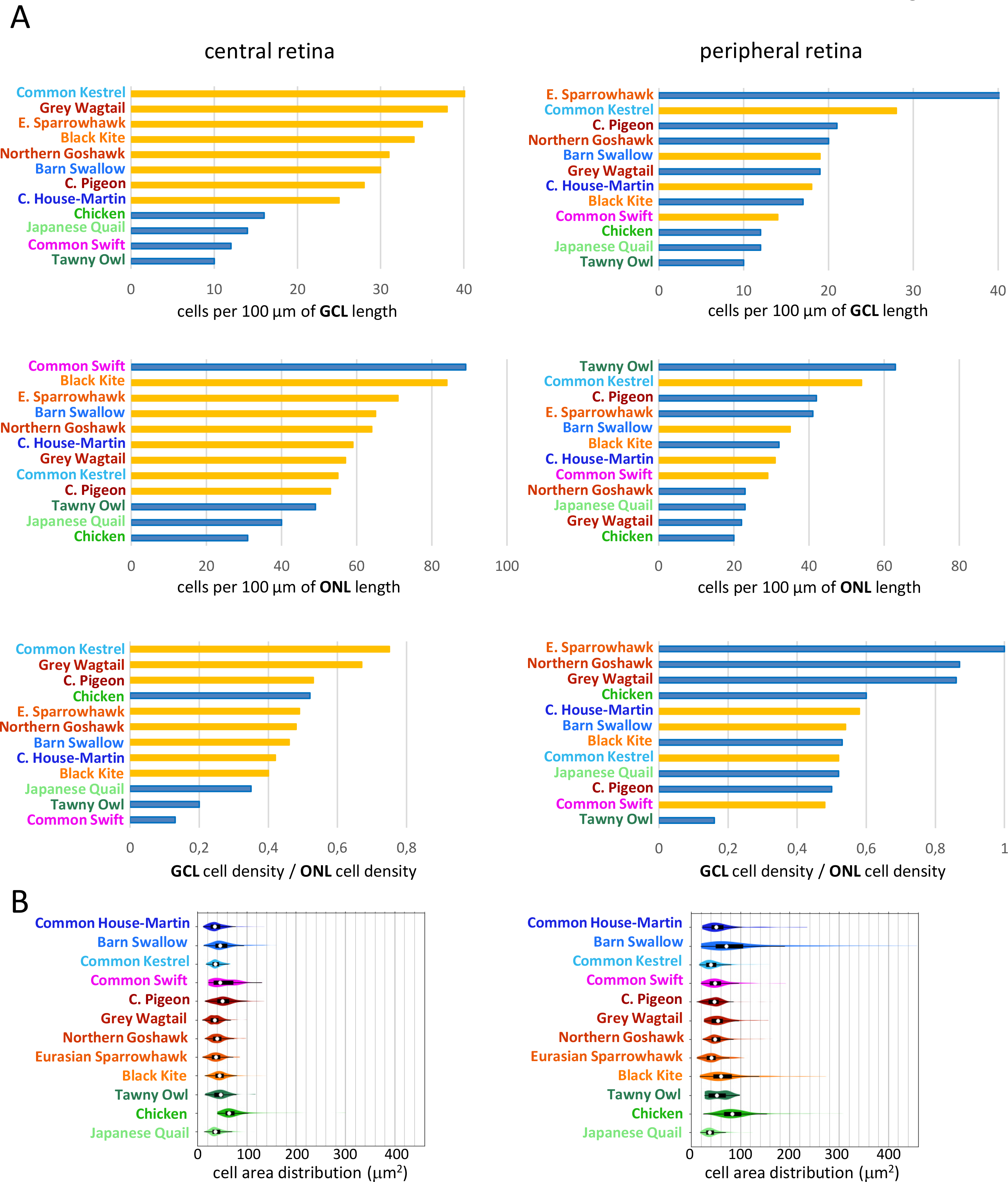
Neuron densities in the retina. (A) Cells in the GCL and ONL were counted on semi-thin cross- sections (1 μm thick) of the foveal and perifoveal areas (∼1 millimeter in length). (A, left panels) Central retina. (A, right panels) Peripheral retina. The orange bars identify the species with a central fovea (left panels) and/or a peripheral fovea (right panels). The species with a central fovea display the highest cell densities in the central GCL. The species with a peripheral fovea are not always the ones that display the highest cell densities in the peripheral GCL. In bifoveate species, cell densities in the GCL are lower in the peripheral fovea than in the central one. The Common Swift has no central fovea, but it displays the highest cell density in the central ONL. Apart from this remarkable exception, cell densities in the ONL are higher in birds with a central fovea. Cell densities are lower in the peripheral ONL than in the central one, except in the Tawny Owl. Cell densities in the peripheral ONL are sometimes lower in species with a peripheral fovea than in species without it. (A, left and right lower panels) In the 12 studied species, the ratios of GCL to ONL cell densities could vary by one order of magnitude. The ratios are < 0.2 in the swift or owl central retinas, while they are equal or close to 1 in the hawk’s peripheral retinas. In 10 out of the 12 central and peripheral foveae examined, the ratios are comprised between 0.4 and 0.6. (B, left panel) Central retina. (B, right panel) Peripheral retina. Each violin plot shows the empirical distribution of the soma surface in the GCL obtained with a gaussian kernel density estimation. On the overlaid box plot, the white dot on each violin plot shows the median, the vertical black bar shows the interquartile range (IQR), whereas the black line shows the range between lower and upper whiskers. Note the high proportion of giant cells (29% ≥ 100 µm^2^) in the peripheral fovea of the Barn Swallow (Figure 2, close-up B).

We asked how this diversity in fovea organization reflects on cell densities in the SGFS. The OT lobes were divided in three sectors: MT1, MT2, MT3 (Figure 1A). In pigeon, the central fovea projects to the lateral part of the OT (Clarke and Whitteridge, 1976) and the MT3 sector encompasses this area. We assumed that the foveal confluence of the OT in the other foveate species is broadly similar to that of the pigeon. For the purpose of this study, there is no need to know precisely where the fovea projects, since RGC densities are much higher across a broad central and peripheral area when there is a central fovea [e.g., the comparison between the pigeon and chicken RGC density maps (Rodrigues et al., 2016)]. Each of the three sector was sectioned perpendicular to the pial surface. Eight 1 µm-thick sections per sector distributed over a thickness of 200-300 µm were selected. On each section the nuclei with visible nucleoli were counted in 10 to 30 contiguous sectors (50 µm x 60 µm) distributed over the surface of the SGFS (Figure 1B, C). We obtained fairly consistent results, despite the fact that we had access to only one specimen for each of the wild species. On the whole, in the two representants of Galloanserae and in the Tawny Owl, the cell densities were lower than in diurnal Neoaves. It was in the bifoveate Common House-Martin, Barn Swallow, and Common Kestrel that cell densities reached the highest levels in all of the three sectors. In the MT3 sector, cell densities were higher in Neoaves than in Galloanserae, with the remarkable exception of the Common Swift and Tawny Owl, strengthening the idea that this sector encompasses the central fovea projection area. Likewise, cell densities in the MT1 and MT2 sectors were the highest in the four species with a peripheral fovea, suggesting that the two sectors share the projection area of this fovea. Cell densities in the MT1 and MT2 sectors were still higher in the five foveate species with no peripheral fovea than in afoveate birds. This is consistent with the fact that RGC densities in the peripheral retinas of foveate birds were higher than in afoveate birds (Figure 3). Unlike in the other Neoaves, in the Tawny Owl, cell densities in the sectors MT1, MT2 and MT3 were at the same low levels as in Galloanserae (Figure 1B, C). Results obtained with this afoveate owl are consistent with the idea that increased cell densities in the largest visual center of the brain are related to the presence of one or two foveae rather than changes that may have occurred when birds split between the Galloanserae and Neoaves.

### 2. The main features of the bird fovea

We found that the cell density in the SGFS of the OT was 2 to 4 times higher in foveate than in afoveate birds. Similar difference was measured between the V1 of primate vs. non-primate mammals (Collins et al., 2010; Rockel et al., 1980; Srinivasan et al., 2015) suggesting that some key features of foveal vision are conserved between diurnal Neoaves and primates. The human fovea is regarded as having the highest spatial resolution of all retinal specializations because of its high density of cones and RGCs and the lowest convergence ratio of cones to RGCs. This allows a concentration of RGCs having small receptive fields, which is presumed to be the substrate for the high acuity pathway. Moreover, in primates, the pit that characterizes the fovea is considered as a way to avoid light scattering by displacing RGCs and interneurons on the rim of the pit as well as to improve the picture that projects on foveal cones by a lens effect.

One common feature of birds with a central fovea is the higher density of RGCs in their central retina and of SGFS neurons in their MT3 sector (Figures 1, 2, 3). As a general trend, densities of photoreceptors, mostly cones, were higher in species with a central fovea. However, there is no central fovea in the Common Swift and the cone density in its central retina is the highest among all species investigated here (Figures 2, 3). Our interspecies comparisons revealed that the relative differences in cell densities between the ganglion cell layer (GCL) and the SGFS varied depending on whether the foveae were deep or shallow (e.g., the comparison between the Common House-Martin and the Northern Goshawk, Figure 4A, B). Likewise, the four species with a peripheral fovea display the highest cells densities in their MT1 and MT2 sectors, while RGC densities within and around their peripheral pit are often lower than in Neoaves with no peripheral fovea (Figures 2, 3, 4). This raises the possibility that a foveal RGC could be connected with a greater number of tectal cells than a RGC outside the fovea, a trend that might be even more pronounced in retinas with a deep fovea.

**FIGURE 4.**
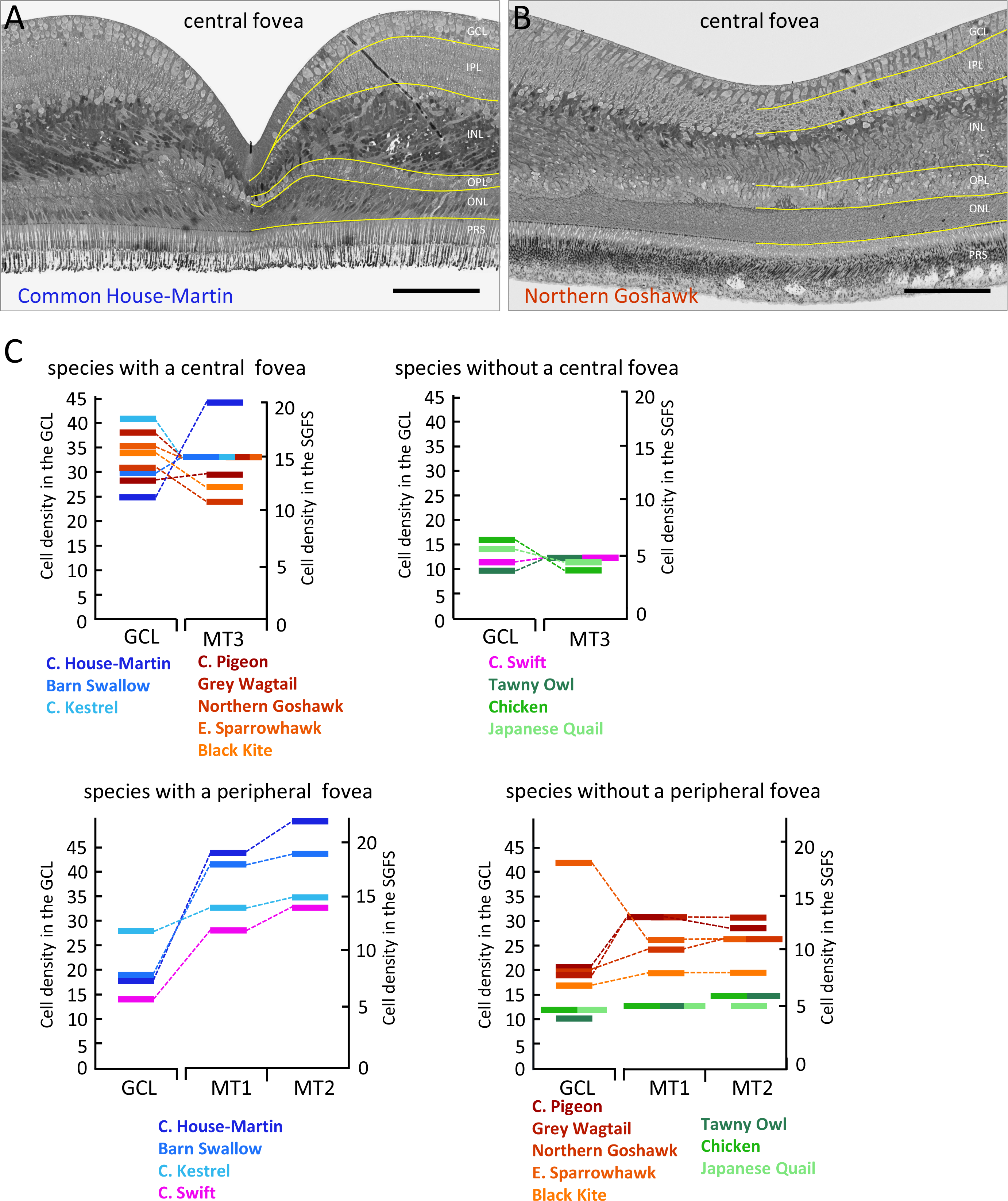
Fovea and cell densities in the GCL and the SGFS. (A, B) Deep vs. shallow foveae. (A) In the deep fovea, the ganglion cell layer (GCL), the inner nuclear layer (INL) and the outer nuclear layer (ONL) are interrupted. The lateral displacement of cells of the INL and ONL leads to the thickening of these layers around the pit. In contrast, the much decreased number of GCL cells within the pit does not result from the displacement of cells on the edge of the pit. (B) In the shallow fovea, there is no layer interruption, but a thinning of the INL. (C) Graphical representations that associate cell densities in the GCL with those in the SGFS (data are from the Figures 1C, 3A). Because of expected variability between individuals of the same species, we compared the cell densities obtained on the same specimens. Meaningful statistics would require a significant number of specimens for each species, which is still a distant goal for wild birds. Species with no fovea display the lowest cell densities in the GCL and in the SGFS. In foveate birds, the relative differences in cell densities between the GCL and the SGFS vary depending on whether the fovea is deep or shallow. The Common House-Martin has the deepest central fovea. Yet, cell density in its central GCL is the lowest among the species with a central fovea, while cell density in the SGFS of its MT3 sector is the highest. There is a similar trend in the peripheral retina. The swallows have deep peripheral foveae and the highest cell densities in the SGFS of their MT1 and MT2 sectors. The Common Kestrel has a shallow peripheral fovea and cell densities in the MT1 and MT2 sectors are comparatively lower, while the cell density in its peripheral GCL is the highest among the species with a peripheral fovea. The Eurasian Sparrowhawk has no peripheral fovea. Yet, the cell density in its peripheral GCL reaches the highest value of all species, while cell densities in the MT1 and MT2 sectors are among the lowest. Scale bars: 100 µm.

### 3. The incipient fovea in the embryonic retina

The pit is the most evident feature of the fovea and its formation likely represents the last step in the morphological development of the fovea, which, in human, is completed one year after birth. In Common Pigeon (herein called ‘pigeon’), we did observe a fully developed pit eight weeks after hatching. Little is known about the developmental processes that precede pit formation. In pigeon, RGC production is completed around embryonic day 12 (E12). At this stage, the retina is about half of the adult size (∼110 mm^2^ at E12 versus ∼250 mm^2^ 1 year post hatching) and the density of RGCs is high across the retina (60-80 cells per 100 µm, Figure 5; (Rodrigues et al., 2016)). Axon growth contributes to the thickening of the nerve fiber layer (NFL) and to the stretching of the retina afterwards. Our data further show that the expansion of the retina from E12 to adulthood led to a ∼5-fold decrease of cell density in the GCL except in a central area of ∼450 µm in diameter (Figure 5). The fact that the NFL was much thinner in the central area suggests that RGC axons growing towards the head of the optic nerve largely avoided this central area that we define as the *incipient fovea*. Its size was basically unchanged from E12 up to 8 weeks post-hatching. However, cell density in the GCL of this incipient fovea was reduced by half 12 days post-hatching (P12), thereby reaching values measured in the fovea at the adult stage (28 cells per 100 µm; Figures 3, 5). It is worth mentioning that the decrease in cell density at P12 correlated with a bending of the retina in the middle of the incipient fovea (Figure 5). In this perspective, it would be interesting to know whether the decrease of cell density initiates the pit formation and whether it is more pronounced in species with a deep fovea. Future studies should also determine whether decreased cell density results from targeted apoptosis.

**FIGURE 5.**
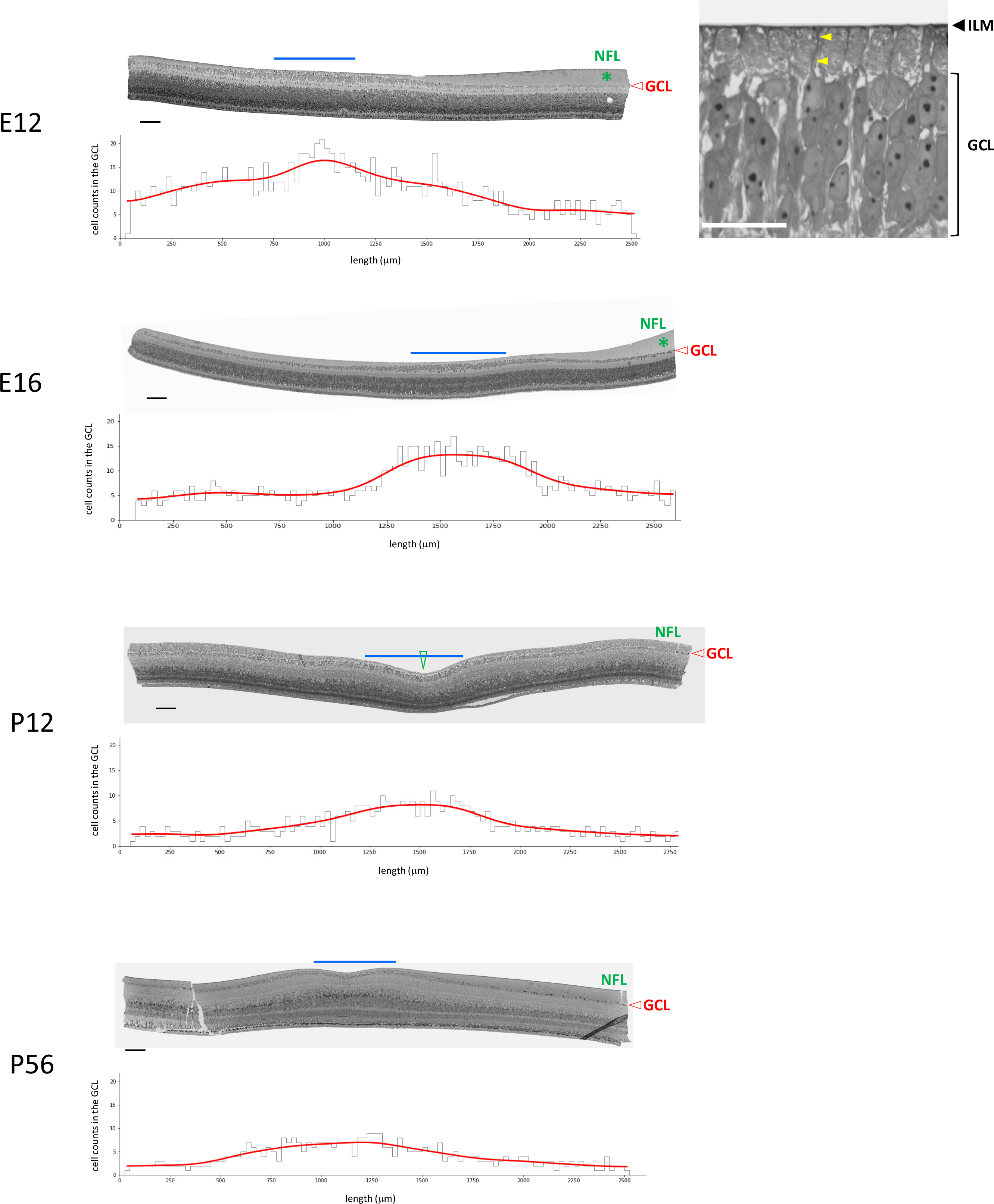
The incipient fovea in the developing pigeon retina. Microphotographs of semi-thin cross- sections of the central retina on the 12th (E12) and 16th (E16) day of embryonic development and on the 12th (P12) and 56th (P56) day of development after birth. The blue line delimits an area where cell density in the ganglion cell layer (GCL) is the highest and the nerve fiber layer (NFL) is the thinnest. Its position and size suggest that it corresponds to the incipient fovea. The NFL, which is made of RGC axons, is absent in the incipient fovea at E12. A close-up of this area shows cells abutting the inner limiting membrane (ILM). The plots show cell densities in the GCL. The gray contour corresponds to the number of cells (on y-axis) in vertical frames partitioning the whole section (on x-axis), each one of them having a width of 25 µm. The red curve is obtained by smoothing the binned values with a gaussian kernel with a standard deviation of 5. At P12, there is a bending of the retina on the middle of the incipient fovea (green arrowhead). Scale bars: 100 µm (E12 to P56), 25 µm (close-up).

### 4. Difference in cell densities between Neoaves and Galloanserae is achieved during embryogenesis

Increased density of tectal neurons in bird species with one or two foveae might represent a breakthrough innovation in visual system evolution. Our cross-species comparison indicates that the pigeon is representative of foveate birds, making comparison between pigeon and chicken an appropriate model to ask how the OT had adapted for receiving higher numbers of retinal inputs. Few days before hatching (E14-E17), the size (length, width, thickness) and the estimated volume of the pigeon OT were, respectively, 1.25 (± 0.1) and ∼2 times smaller than in chicken (Figure 6A, D). In pigeon, the SGFS layers receive three times more RGC axons than those of chicken at E17 (Figure 6B; Rodrigues et al. (2016)). The stratum opticum (SO) is the most superficial layer of the tectum and it is formed by RGC axons of the optic tract. At E14, the SO was thicker in pigeon than in chicken all along the medio-lateral or the rostro-caudal axis (Figure 6D and data not shown). From E14 through E17 (P0), we noted several histoarchitecture differences between the pigeon and the chicken. At E14, in the chicken OT, LaVail and Cowan (1971a) identified on 20 µm thick paraffin sections 10 layers, numbered i-x from the ependymal surface. On 1 µm semi-thin plastic sections, the cellular layers iv, vi, viii and x were clearly visible in pigeon, whereas they were blurry in chicken, because of the lower cell density (Figure 6D). Cell counting revealed that neuron densities in the layers vi to x were 2 to 4 times higher in pigeon than in chicken at E14 (Figure 6D, E, F). At this stage, the total number of tectal cells was actually the same in pigeon and chicken (Table S2). However, in pigeon, cells were packed in a volume twice smaller. Difference in neuron densities between the pigeon and chicken outer layers exceeded those measured in isotropic suspensions of cell nuclei, reflecting non-uniform scaling with comparatively higher cell densities in the outer laminae where RGC axons terminate. Likewise, at the adult stage, the 2.5 (MT1), 2.2 (MT2), 2.8 (MT3) times higher cell densities in pigeon compared to chicken (Figure 1C) mainly concern cells localized in the superficial layers of the SGFS (Figure 6C). Taken together, our data indicate that the difference in cell density between the pigeon and the chicken were already established at E14.

**FIGURE 6.**
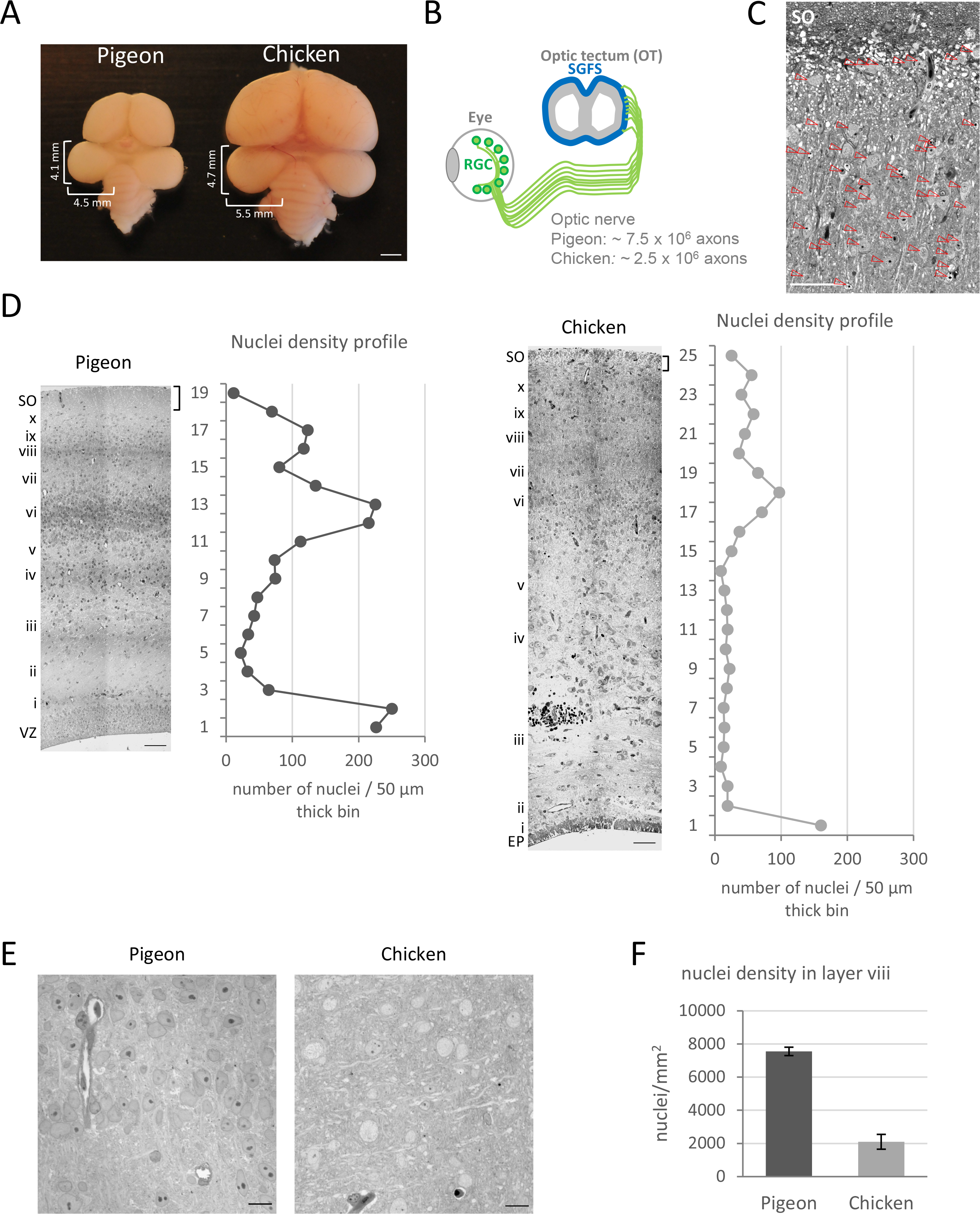
Cell density patterns in the developing pigeon and chicken OTs. (A) Pigeon and chicken brain at E17. (B) RGCs make direct connections with SGFS neurons. The number of axons in the optic nerve is ∼3 times higher in pigeon than in chicken (Rodrigues et al., 2016). (C) Semi-thin cross-section through the outer layers of the SGFS in the MT3 sector of an adult (P70) pigeon. Nuclei are indicated by arrowheads. (D, left panels) Semi-thin cross sections of pigeon or chicken OT at E14. The layers recognizable at this stage are numbered in sequence from the ventricular zone (VZ, in pigeon) or the ependymal surface (EP, in chicken) and the outward according to the convention established by LaVail and Cowan (1971a). (D, right panels) The curves show the number of cells (on x-axis) in horizontal frames partitioning the whole section (on y-axis). Similar cell density profiles were obtained on OT sections from 3 embryos for each species. (E) Close-up pictures of the layer viii. (F) The histogram represents mean ± s.d., calculated in ∼50 frames on OT sections from 3 embryos for each species. Scale bars: 2 mm (A), 40 µm (C), 50 µm (D), 10 µm (E).

### 5. The onset of neurogenesis in the pigeon and chicken optic tectum

Like other altricial birds, the pigeon hatches with closed eyes. We wondered whether the prolonged OT development after birth (Bagnoli et al., 1987; Manns and Gunturkun, 1997) could result from delayed neurogenesis. To answer this point, we first compared how neurogenesis unfolds in the pigeon and chicken OTs at early stages of embryonic development. Measuring the sizes, the number of cells and the number of cell division per day indicated that the pigeon and chick OTs grew and expanded at a similar pace from E3 to E6 (Figure 7A; Table S2). Monitoring Ngn2 expression revealed that neurogenesis peaked between E6 and E7 in both species (Figure 7C). NeuroD4 (formerly named NeuroM) marks OT cells at the time of birth and when they are located in a zone immediately superficial to the neuroepithelium (Roztocil et al., 1997). NeuroD4 transcript accumulated according to similar kinetics in pigeon and chicken suggesting that cell birth and the onset of neuronal differentiation follow the same trend (Figure 7C). This is also reflected in the fact that axon growth was detected from E5 in both species (Figure 7D). Until E6, the pigeon OT had the form of a thin-walled vesicle consisting of an ependymal zone layered with neural epithelial cells and newborn neurons defining the ventricular zone (VZ) and an outer layer formed by a meshwork of radial fibers (Figure 7B). At E6, we noticed that cells had already started to migrate between glial fibers in chicken, whereas that was not yet the case in pigeon. At this stage, we had no evidence that the onset of neurogenesis was delayed in the pigeon OT, in contrast to what happens in the retina (Rodrigues et al., 2016).

**FIGURE 7.**
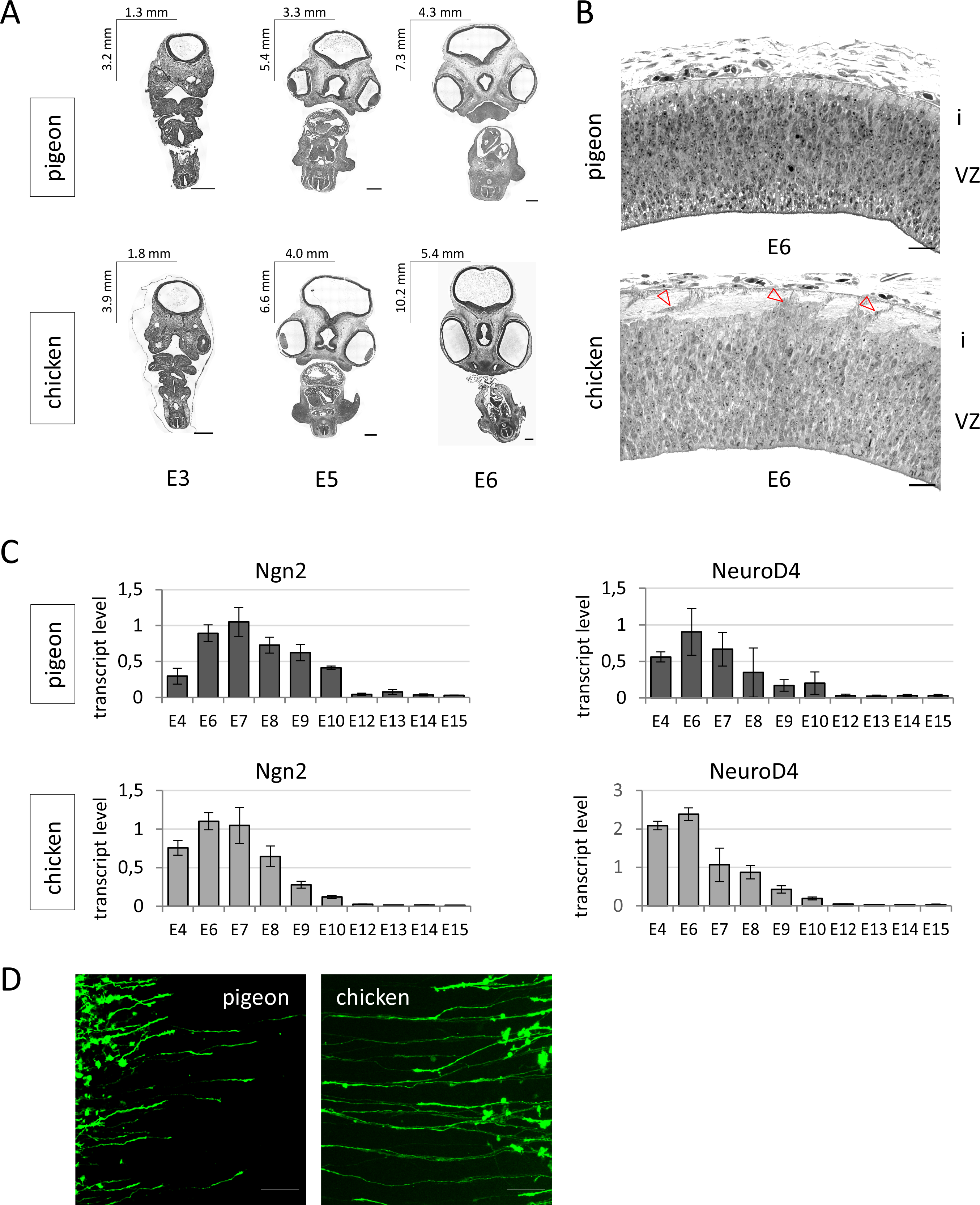
Early development stages of the pigeon and chicken OT. (A) Frontal sections through the whole embryo at the level of the midbrain. The OT vesicles are on the top of the pictures. The length and width of tissue section are indicated in the upper-left corner (B) Semi-thin cross- sections of the central portion of the OT, i.e., midway between its dorsomedial and ventrolateral borders. Superficial to the VZ lies the outer glial fibers (layer i). The arrowheads in chicken OT point to cells positioned between glial fibers. (C) Accumulation of Ngn2 and NeuroD4 (NeuroM) transcripts measured by RT-qPCR analysis. At each developmental stage, data are from biological triplicates and presented as the mean ± s.d. (D) Pigeon and chicken OT were electroporated in ovo with a Hes5.3-GFP reporter plasmid at E4.5. Labelled axons were visualized 24 hours later. Scale bars: 500 µm (A), 25 µm (B), 50 µm (B).

### 6. Interspecies divergence of developmental trajectories

On the 7^th^ day of embryonic development, the pigeon and chicken OTs displayed similar radial glia fascicles consisting of equally spaced parallel thick rows of glial fibers abutting the VZ (Figure 8A, B; Gray and Sanes (1991)). However, whereas in chicken cells arranged in single files migrate along glial fibers to form the tectal plate, in pigeon no cell filled the fascicle interspaces and the tectal plate did not form. The deepening of the groove separating the right and left optic lobes proceeded at a fast pace in chicken, whereas the two lobes did not separate in pigeon. In chicken, most cells were born at E7 (Figure 10A, Table S2) and the lamination led to a rapid thickening of the OT. In contrast, in pigeon, there was no cell migration out of the VZ between E7 and E9 (Figure 8A, B). To investigate further the significance of these striking cross-species differences, we compared in pigeon the development periods that extended from E6 to E7 and from E7 to E9 (Figure 9). The number of cells doubled during each of these periods. The 3D reconstruction of the OT with 10 µm thick serial sections showed that the tissue volume increased 2-folds from E6 to E7 and 2.5-folds from E7 to E9. However, the growth pattern of the tissue had drastically changed over the E6 to E9 period of development. Between E6 to E7, the tissue expanded tangentially and its length along the rostro-caudal and cross axis increased, while the thickness of the tectal tissue and the number of cells in radial columns increased slightly. This stands in sharp contrast to what happened from E7 to E9. During this latter period, there was no tangential growth along the rostro-caudal axis, whereas there was a marked thickening of the tissue and the number of cells in radial columns encompassing the VZ increased by ∼60% (Figure 9). Our histological and morphometric analysis suggest that late cell division in the pigeon happens in a cellular environment different to that prevailing up to E7. Late cell proliferation in a VZ expanding radially - i.e., almost exclusively along its apicobasal axis - led to an increase in the number of cells in radial columns, thereby setting the condition for an increase of cell density. This model of tissue morphogenesis raises a number of questions about the status of cells stuck in the thickened VZ abutting the glial scaffold (Figure 8C).

**FIGURE 8.**
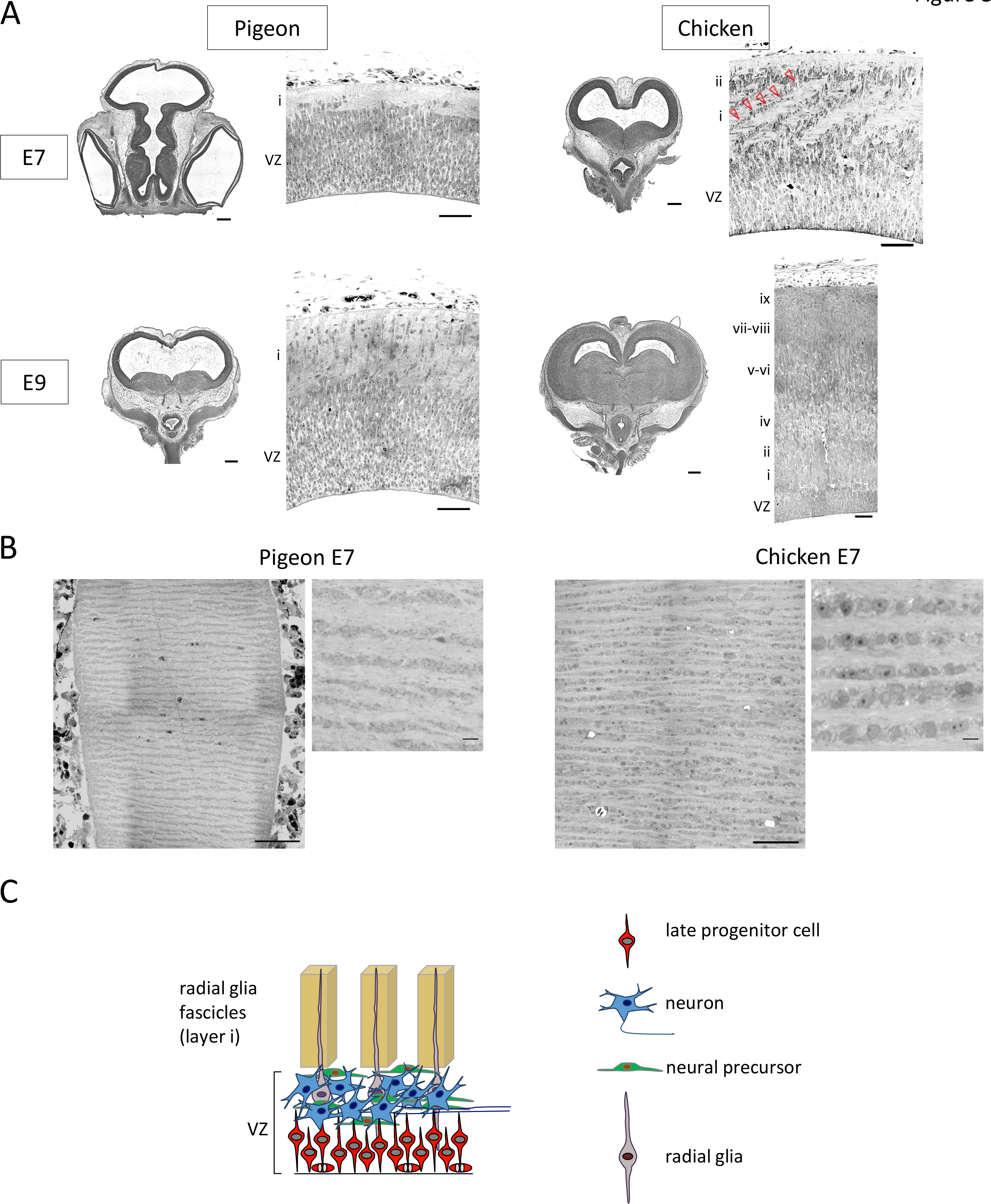
Divergent development of the OT between pigeon and chicken. (A, left panels) Low magnification pictures of frontal sections through the OT. (A, right panels) Semi-thin cross-sections of the central OT, i.e., midway between its dorsomedial and ventrolateral borders. At E7, the thicknesses of the VZ are the same in pigeon and chicken. From E7 to E9, VZ thickness increases by ∼40% in pigeon, whereas it decreases by ∼45% in chicken. At E9, the VZ is ∼2.5 times thicker in pigeon than in chicken. In chicken, at E7, cells arranged in single files (arrowheads) migrate along glial fibers (layer i), heading towards the outer layer ii; at E9, OT thickness has increased and 9 layers (i-ix) can be identified. In pigeon, at E7 and E9, the two lobes do not yet separate. At E9, rare cells are scattered in the layer i, but none of them did cross this layer. (B) Semi-thin sections cut tangentially through the fiber layer i of pigeon and chicken OT at E7. In chicken, rows of cell-rich and fiber-rich stripes alternate, whereas in pigeon, there is no cell between the fiber-rich stripes. (C) Organization of the pigeon OT at E9. This preliminary view is based on data presented in the panels A, B and in the Figures 10D and 11C. Scale bars: 500 µm (A, heads), 50 µm (A, OT sections; B), 5 µm (B, close-up pictures).

**FIGURE 9.**
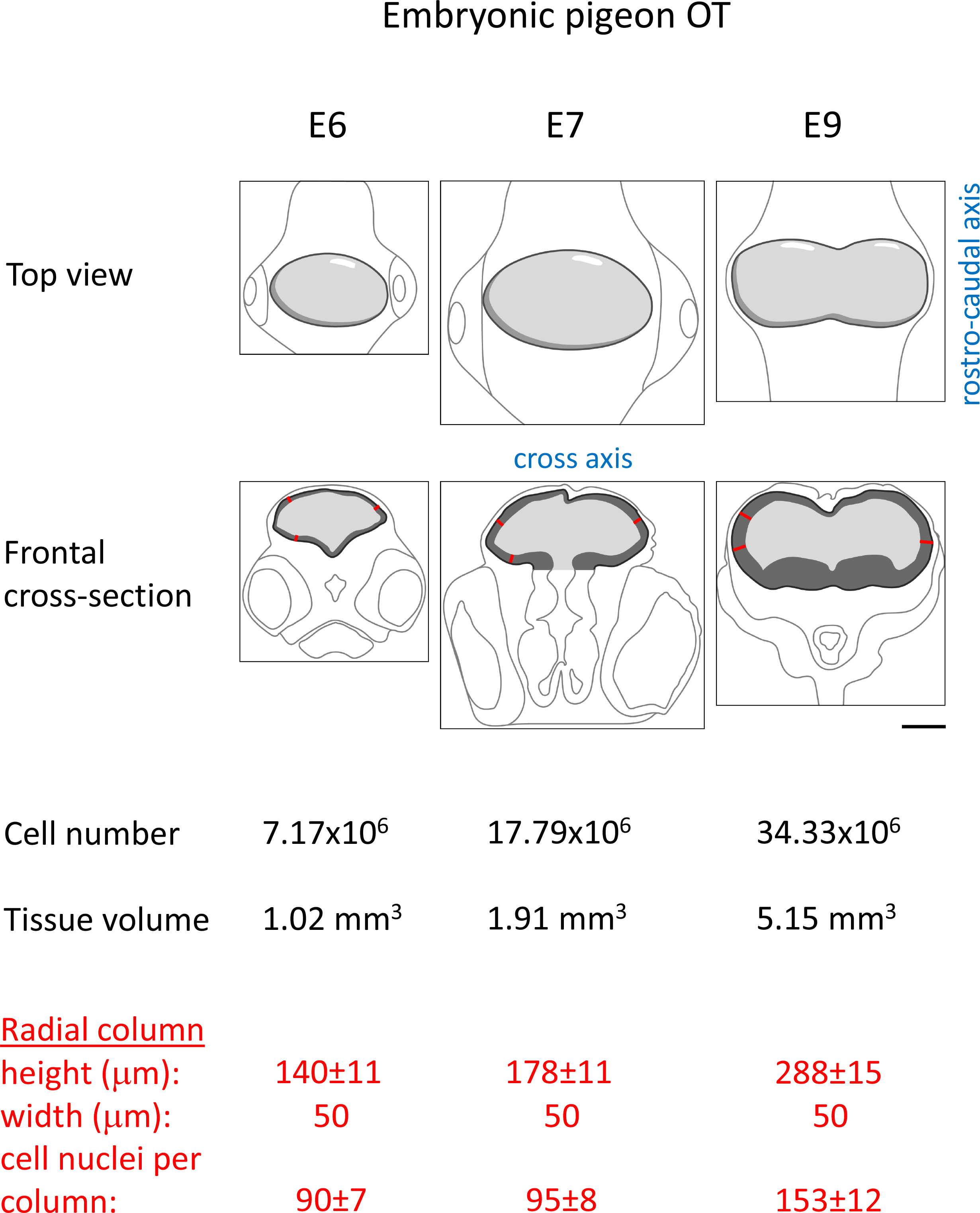
Change of the growth pattern of the pigeon OT. Cell number doubles from E6 to E7 and again from E7 to E9. The increase in OT tissue volume from E6 to E7 mainly reflects tangential expansion. In contrast, the OT does not expand along its rostro-caudal axis from E7 to E9 and the increase in OT tissue volume mainly reflects increased radial thickness. Cell number data were obtained from isotropic suspensions of cell nuclei (Table S2). Tissue volume data were obtained by 3D reconstruction of the OT with serial cross-sections. The height of the radial column corresponds to the distance (thickness) between the ventricular and the outer surfaces measured on ventrolateral sections. Cells per radial column correspond to the number of nuclei counted on semi-thin cross-sections in 50 µm wide sectors. Scale bar: 1 mm.

**FIGURE 10.**
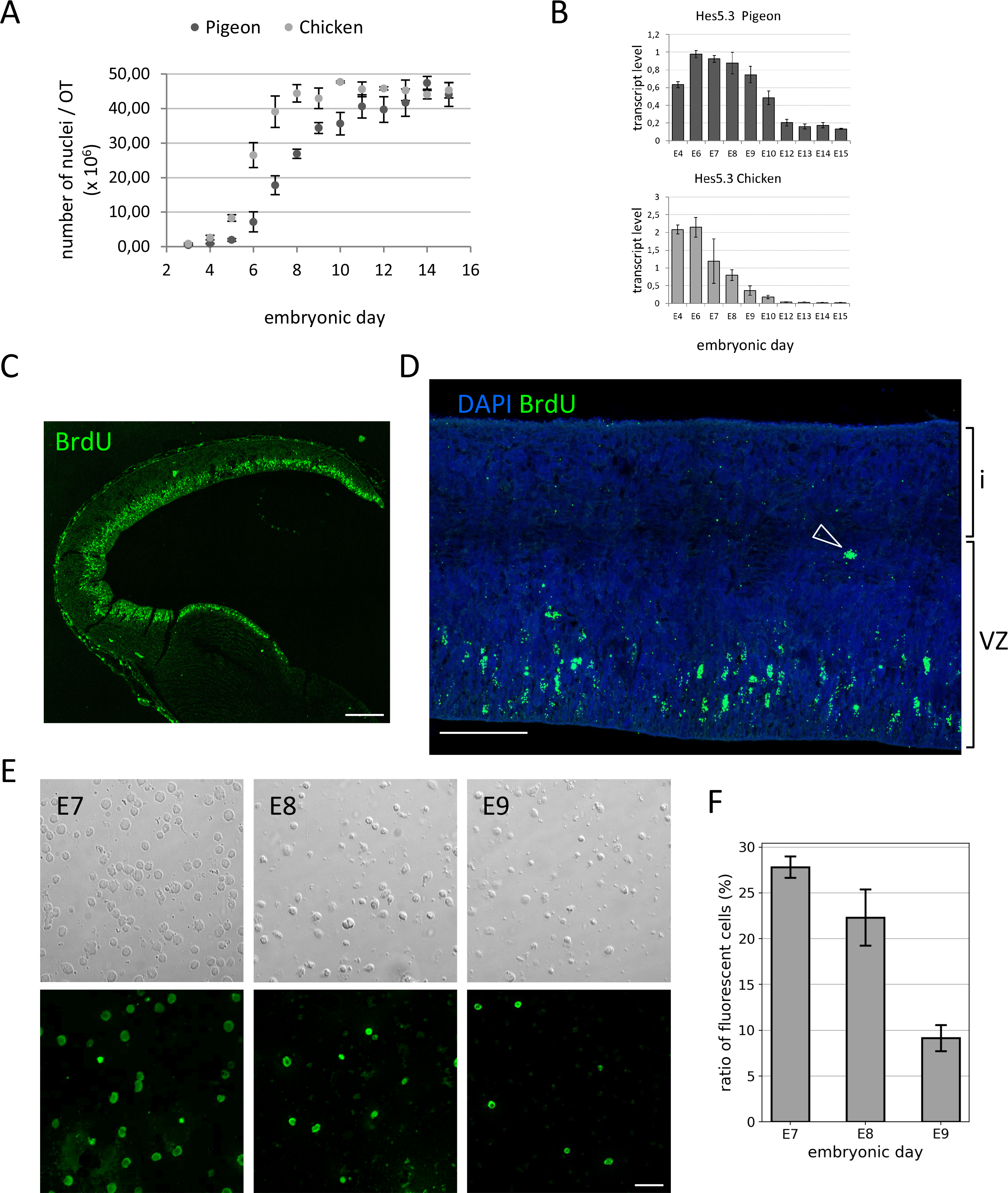
Late cell proliferation in the pigeon OT. (A) Cell counts obtained with isotropic suspensions of OT cell nuclei. Each point of the curves is the mean ± s.d. of counts from biological triplicate (Table S2). (B) Accumulation of Hes5.3 transcripts measured by RT-qPCR analysis. At each developmental stage, data are from biological triplicates and presented as mean ± s.d. (C-F) BrdU pulse-labelling experiments. OTs were dissected at E8 (C) or E9 (D) and incubated as explants with BrdU for 1 hour. After a 1 hour chase, the tissues were fixed and embedded in paraffin. Sections were processed for immunohistochemistry (C, D) and counterstained with DAPI (D). (D) > 95% of labelled cells are localized on the apical side of the VZ. Rare cells (e.g., the one indicated by the arrowhead) are detected on the basal side of the VZ. (E, F) OTs were dissected at E7, E8 or E9 and incubated as explants with BrdU for 19 hours. Dissociated cells were processed for immunohistochemistry. (E) Phase-contrast and fluorescence images are shown. (F) Percentage of BrdU-labelled cells. Data are from biological triplicate and are presented as mean ± s.d. Scale bars: 200 µm (C), 50 µm (D), 20 µm (E).

### 7. Late progenitor cells divide in a radially expanding ventricular zone

The length and width of the pigeon embryos are ∼20% smaller than the chicken embryos and the OTs follow this general trend (Figures 7A, 8A). Recalling that the volume of a cube with a side length of 0.8 mm is half the volume of a cube with a side length of 1 mm, the volume of the pigeon OT at E7 is about half the volume of the chicken OT. Likewise, the pigeon OT contains about half as many cells as the chicken OT [∼18 x 10^6^ cells in ∼1.9 mm^3^ for the pigeon vs. ∼39 x 10^6^ cells in ∼4.2 mm^3^ for the chicken (Figures 9, 10A)]. In chicken, the number of tectal cells was close to the maximum level at E7, whereas, in pigeon, at this stage, cell number was only at ∼50% of the maximum level that was reached around E11 (Figure 10A). When E8 or E9 pigeon OT explants were pulse-labelled with BrdU for 1 hour, labelled cells were abundant all over the dorsoventral surface of the tectum (Figure 10C). More than 95% of labelled cells were localized on the apical side of the VZ, whereas very few were found on the basal side abutting radial glia (Figure 10D). Cells of the VZ were tightly packed and their distribution looked quite homogeneous (Figure 8A). To estimate the fraction of cells that still proliferate, E7, E8 and E9 OT explants were incubated with BrdU for 19 hours. Dissociated cells were plated on slides and processed for immunodetection (Figure 10E, F). The proportions of labelled cells suggest that a pool of progenitor cells corresponding to ∼28% of the total number of cells divided between E7 and E8, and that another pool of similar size divided again between E8 and E9. The much decreased number of labelled cells detected between E9 and E10 suggests that the period of prolonged cell proliferation was completed shortly after E10. The data of this experiment allow to calculate a ∼1.7-fold increase in cell number between E7 and E10. A similar increase was measured with the isotropic suspensions of cell nuclei (Figure 10A). Hes5.3 is expressed in progenitor cells during the penultimate and last cell cycle (Chiodini et al., 2013). In pigeon, Hes5.3 transcript level remained close to its peak value up to E9, whereas it rapidly decreased in chicken after E6 (Figure 10B).

### 8. In pigeon, different genes regulate proliferation of the early and of the late progenitor cells

In pigeon, the late cell proliferation (E7 to E10) occurred in a cellular context different from the one that prevailed up to E7 (Figures 8, 9). To analyze molecular features of OT cells, we compared the diversity of the pigeon and chicken RNA-Seq transcriptomes at E6, E8 and E10. In chicken, we anticipated that DNA replication, cell cycle progression and mitosis would be one of the major source of transcriptional variations between E6 and E8 because the large majority of cells have stopped to divide at E7 (Figure 10A; LaVail and Cowan (1971b). Indeed, 97 genes involved in cell proliferation were downregulated in chicken during this period (Figure 11A). In pigeon, the situation was highly contrasted and we split the set of 97 genes into three subsets: 1) 62 genes displayed no significant change of expression between E6 and E10, likely reflecting the prolonged period of cell proliferation; 2) 22 genes that initiate DNA replication like those encoding the ORC1 and MCM5 subunits or those representing G1 and G1/S checkpoint like CCND1, CDKN3, CDCA7, CKS2 were downregulated in pigeon between E6 and E8, much like in chicken; 3) quite surprisingly, 13 genes were specifically upregulated in pigeon between E6 and E8, and, for some of them, between E8 and E10. For example, the gene POLE that encodes the central catalytic subunit of the DNA polymerase epsilon was strongly upregulated during the two periods in pigeon (+3.24 E6-E8; +1.92 E8-E10 in log2), whereas it was downregulated in chicken (-1.43 E6-E8; -0.50 E8-E10 in log2). On the other side, ORC1 was strongly downregulated both in pigeon (-4,88 in log2) and in chicken (-2,53 in log2) between E6 and E8, whereas the expression of ORC2-6 did not vary in pigeon during this period. ORC1-6 binds to origins of replication as a ring-shaped heterohexamer and the subunit ratios could vary (Shibata et al., 2016). Another major interspecies difference was that genes representing G2/M checkpoint like CCNB2, TICRR or key steps in cellular response to DNA damage like ATR, FANCL, MCPH1 (microcephalin), NDE1 and TSN (translin) were strongly activated in pigeon between E6 and E8 (e.g., +3,77 for TSN; +3.39 for MCPH1 in log2), whereas they were downregulated in chicken (-1.10 for TSN; -2.17 for MCPH1 in log2). Mutation or ablation of ATR, MCPH1 or NDE1 in mice induce brain malformations (Feng and Walsh, 2004; Gruber et al., 2011; Lang et al., 2016; Lee et al., 2012). In pigeon, the vast majority of late progenitors generate neural cells of the SGFS (see section *Fate and positioning of cells produced by the late pool of progenitors*). The late upregulation of genes controlling proliferation (and, for some of them, genome stability) could be a feature of progenitors that generate neural cells of the SGFS. However, the fact that in chicken these genes were already downregulated when progenitors that generate neural cells of the upper layers of the OT were still proliferating (Figure 11A; LaVail and Cowan (1971b)) argues against this idea. We suggest that these genes are specifically activated in pigeon in response to extrinsic conditions that may prevail in the VZ from E7 to E10 (Figure 8C; e.g., the environmental stress resulting from neurons-progenitor cell proximity). Gain- and loss- of function experiments will be needed in future studies to determine the role of genes differentially expressed in pigeon and chicken. However, the rather poor annotation of the pigeon genome and the current lack of breeding facilities for raising genetically modified hatchlings hinder progress towards this goal.

**FIGURE 11.**
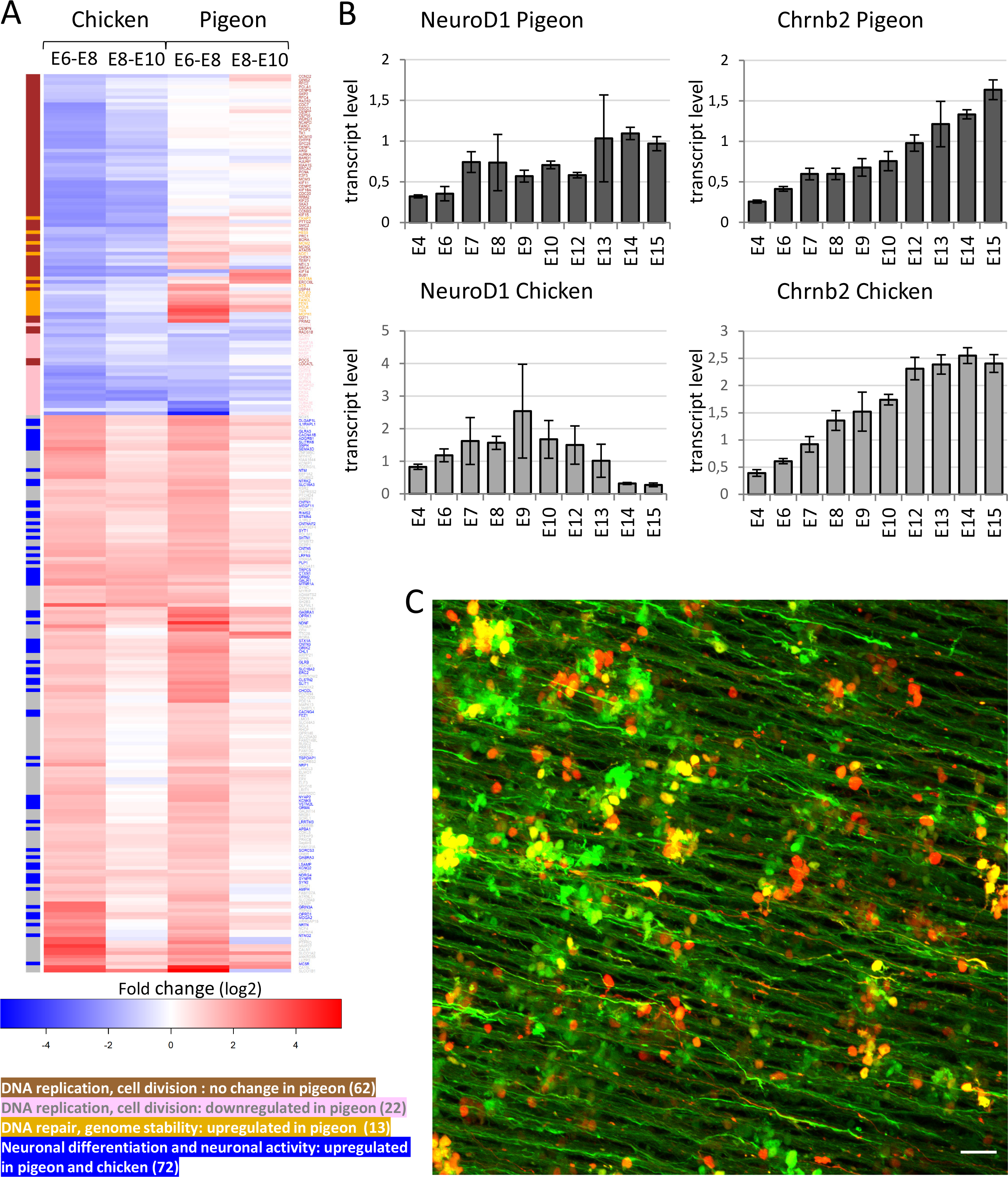
Transcriptomic analysis of the pigeon and chicken OTs. (A, upper panel) Heat maps generated from RNA-Seq data showing clustered genes regulated during the two periods between E6 and E10. (A, lower panel) In chicken, 97 genes known to regulate DNA replication and cell division are downregulated between E6 and E10. In pigeon, 62 orthologs (gene symbol in brown) show no significant change in expression, 22 orthologs (gene symbol in pink) are downregulated, 13 orthologs (gene symbol in orange) are upregulated between E6 and E10. Out of the 166 genes upregulated in both pigeon and chicken during the period between E6 and E8, 72 genes (gene symbol in blue) are known to be involved in neuron differentiation and neuron activity. 94 genes (gene symbol in grey) have not been submitted for gene ontology analysis. (B) Accumulation of NeuroD1 and Chrnb2 transcripts measured by RT-qPCR analysis. At each developmental stage, data are from biological triplicates and presented as mean ± s.d. (C) Axons of neurons residing in the VZ. The pigeon OT was co-electroporated in ovo with Hes5.3-GFP and CMV-RFP reporter plasmids at E4. Fluorescent axons and somas were visualized by confocal microscopy at E7. The image plane is normal to the apico-basal axis and situated at the level where radial glial fascicles abut the VZ as evidenced by scattered cell bodies. Parallel bundles of fluorescent axons are at intervals corresponding to the distance between the radial fascicles (5-10 µm; Figure 8B). The vast majority of axons are exclusively labeled with GFP, consistent with the fact that Hes5.3 is preferentially expressed in new born neurons. Scale bar: 20 µm.

### 9. Neurons differentiate while they remain stuck in the ventricular zone

Out of the 166 genes that were upregulated in both the pigeon and chicken OT between E6 and E8 and between E8 and E10, 72 are involved in neuronal differentiation and neuronal activity. This set includes 27 genes that regulate axonal and dendritic outgrowth, 21 genes that encode receptors and ion channels and 17 genes involved in synaptogenesis and synaptic transmission (Figure 11A). It is important to remember that genes regulating neuron differentiation in pigeon from E6 to E8 were exclusively expressed by cells residing in the VZ (Figure 8A). Gene expression profiling in the developing brain of birds is still in its infancy. However, it is well established that cholinergic circuits are involved in the visual information processing (reviewed in Knudsen (2011)) and that both SGFS neurons and axon terminals of RGC afferents express nicotinic cholinergic receptors (King, 1990; Matter et al., 1990; Titmus et al., 1999). The neuronal acetylcholine receptor subunit beta-2 (Chrnb2, formerly named non-alpha nAChR) is predominantly expressed in the SGFS (Matter et al., 1990). Chrnb2 transcript steadily accumulated between E4 and E12 both in pigeon and chicken (Figure 11B). However, while transcript level reached a plateau in chicken at E12, it was still increasing in pigeon at E15. NeuroD1 is a regulator of neuron differentiation in the avian nervous system (Matter-Sadzinski et al., 2001; Roztocil et al., 1997) and the expression patterns of this gene were similar in pigeon and chicken until E12. Then, NeuroD1 was rapidly downregulated in chicken, whereas, in pigeon, expression was maintained at high levels at least until E15 (Figure 11B), indicating that neuron production was extended over several days. Axon growth in the pigeon OT was first detected at E5 (Figure 7D). When Hes5.3-GFP and CMV-RFP reporters were co-electroporated in the OT at E4, regularly spaced parallel bundles of GFP-positive axons were detected at E7 (Figure 11C). Bundle thickness (5-10 µm) suggests that axons were growing between the rows of glial fibers (Figure 8B) but close to the VZ as evidenced by the presence of GFP- and/or RFP-labelled cells. Taken together, our data suggest that neural differentiation unfolded at a similar pace in pigeon and chicken. The fact that in pigeon, neurons differentiated while their somas resided in the VZ is likely to challenge our current understanding of how a laminated brain structure develops.

### 10. Unlocking cell migration

In pigeon, few cells had moved in the interspaces between radial glia fascicles until E9, while at E10, cells migrated through the glial scaffold (layer i) for establishing the outer layers (Figure 12D), a pattern resembling the one in chicken at E7 (Figure 8A). Cell migration out of the VZ coincided with tissue folding that led to the deepening of the median tectal groove and the separation of the two lobes three days later than in chicken (Figure 12A). About half of the cells have already left the VZ at E10 and the proportion of cells in the layers i and ii were the same in the dorsomedial and ventrolateral sectors (Figure 12B). This is different from the situation in chicken where the retarded dorsomedial sector is ∼2 days behind the development of the ventrolateral sector (LaVail and Cowan, 1971b).

**FIGURE 12.**
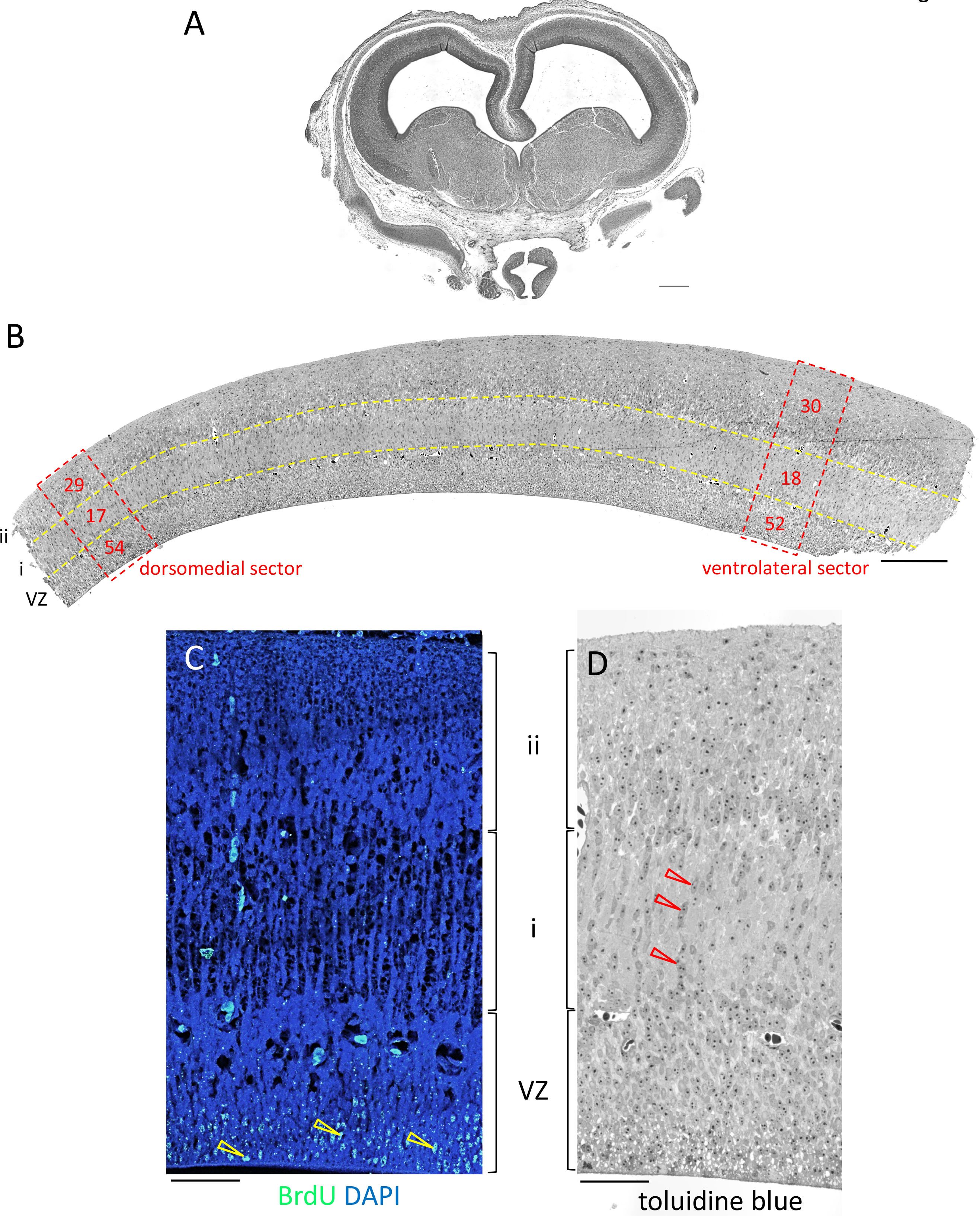
Unlocking cell migration in pigeon. (A) Low magnification picture of a frontal section through the pigeon OT at E10. The separation of the two lobes is underway. (B) Semi-thin section along the dorsoventral axis of the OT at E10. The values in the frames correspond to the proportion of cells (%) in the three layers (VZ, i, ii) delimited by the dashed lines. (C) BrdU was injected in ovo at E8. The OT was isolated at E10 and sections were processed for immunohistochemistry and counterstained with DAPI. Labelled cells (yellow arrowheads) are localized on the apical side of the VZ. (D) Semi-thin section of the central OT at E10. Numerous cells migrate between meshwork of radial fibers in the layer i (red arrowheads). Scale bars: 500 µm (A), 200 µm (B), 50 µm (C, D).

### 11. Cell migration in pigeon and chicken are regulated by different sets of genes

In the chicken OT, neuronal precursors migrate and establish layers between E6 and E8 (Gray and Sanes, 1991; LaVail and Cowan, 1971b), whereas, in pigeon, neurons get stuck in the VZ up to E9. We asked how this spectacular interspecies difference was related to changes in gene regulation. Evsyukova et al. (2013) have compiled a comprehensive list of 165 genes that regulate cortical neuronal migration in mammals. Our transcriptomic data show that 46 genes from this list were up or downregulated in chicken between E6 and E8, whereas their expression did not significantly vary in pigeon. During the same period, 17 genes were up or downregulated in both species (Figure 13A). These two sets of genes encode receptors, secreted molecules, substrate- or membrane-bound molecules or transcription factors that might be involved in cell migration and/or axon growth and guidance. In pigeon, neurons extended axons while their somas still resided in the VZ (Figure 11C), suggesting that the 17 genes differentially expressed between E6 and E8 in both pigeon and chicken (e.g., SLIT1, MAP2, Figure 13A) are likely to be involved in axon growth. One striking feature of our data is that none of the 46 genes differentially expressed in chicken between E6 and E8 were differentially expressed in pigeon between E8 and E10 - i.e., during the period when neurons migrate out of the VZ (Figures 12, 13A). This may reflect the fact that migrating cells were neural precursors in chicken, whereas they were neurons in pigeon. In the developing mouse telencephalon, Nkx2-1 represses the expression of Neuropilin-2 (Nrp2) and the downregulation of Nkx2-1 is a necessary event for the migration of interneurons to the cortex (Nobrega-Pereira et al., 2008). In the same way, Nrp1 and Nrp2 were upregulated while Nkx2- 1 was strongly downregulated in the chicken OT (+2.11 and +2.55 in log2 for Nrp1 and Nrp2; -7.03 in log2 for Nkx2-1; Figure 13A). However, these genes were not differentially expressed in pigeon. The upregulation of the β1 integrin, ITGB1, and of the ligands and activators of integrin, CXCL12, in pigeon are clues pointing to neuron migration dependent on the integrin signaling pathway (Figure 13A, see also Galileo et al. (1992)). The upregulation of the GDNF family receptor alpha GFRA1 and the downregulation of the adhesion molecule NCAM1 in pigeon between E8 and E10 (Figure 13A) are consistent with this idea. In this context, it is worth mentioning that by impeding the function of NCAM, GFRA1 promotes the migration of Purkinje cells in the developing mouse cerebellum (Sergaki and Ibanez, 2017).

**FIGURE 13.**
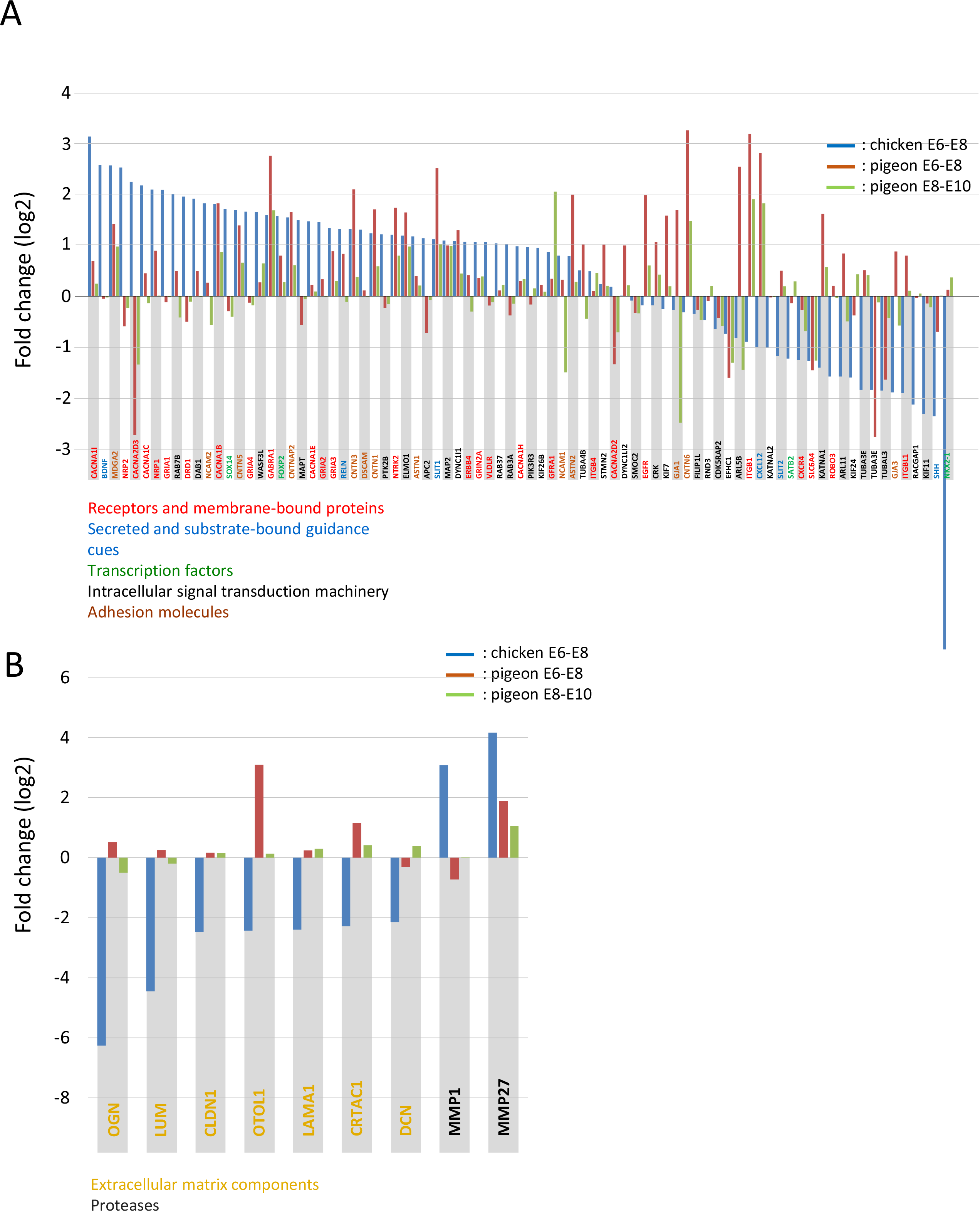
Genes regulating cell migration are differentially expressed in pigeon and chicken OTs. (A, B) Bar graph generated from RNA-Seq data. Positive or negative fold changes in log2 are presented (on y-axis). (A) The graph shows a selection of 83 genes known to regulate cortical neuronal migration and axon growth. 46 genes are up or downregulated (>1 or <-1 in log2) in chicken between E6 and E8. None of them are differentially expressed in pigeon between E6 and E8 or between E8 and E10. 17 genes were up- or downregulated in both species between E6 and E8. (B) The graph shows the regulation of genes encoding extracellular matrix components and matrix metalloproteinases.

We have been intrigued to find that genes encoding extracellular matrix structural and non-structural constituents [Decorin (DCN), Lumican (LUM), Cartilage acidic protein 1 (CRTAC1) Osteoglycin (OGN), Otolin (OTOL1)], components mediating cell-matrix interactions [Laminin (LAMA1)] and tight junctions’ proteins [claudins (CLDN1)] were strongly downregulated at the onset of cell migration in chicken, while their expression did not vary in pigeon, except for OTOL1 and CRTAC1 which were upregulated (Figure 13B). DCN, LUM, CRTAC1, OGN, CLDN1 have the capacity to impede cell movement and could, for some of them, inhibit metastasis (Appunni et al., 2019; Deckx et al., 2016; Hagen, 2017; Karamanou et al., 2021; Yang et al., 2021). The idea that these molecules may contribute, in different ways, to pose physical barriers that prevent cells to migrate out of the VZ is strengthened by the fact that genes encoding the matrix metalloproteases MMP1 and MMP27 were strongly upregulated in chicken, whereas they were upregulated weakly or not at all in pigeon (Figure 13B). The spectacular cross-species differences in the patterns of expression of genes that regulate cell migration could be one crucial feature of the divergence in OT development between pigeon and chicken and, more broadly, between diurnal Neoaves and Galloanserae.

### 12. Fate and positioning of cells produced by the late pool of progenitors

Cross-species differences in the timing and regulation of cell migration raise questions in the context of the lamination of the optic tectum and, in particular, of its ’inside-out’ type of developmental gradient. In this perspective, we wanted to determine the fate and the positioning of cells produced by the late pool of progenitors in pigeon. BrdU was injected in ovo at E8 and tissue sections were probed at E10 or E12 (Figures 12C, 14A, B). At E12, all labelled cells have migrated in the most superficial layers of the tectum, indicating that progenitors dividing beyond E8 were in their vast majority committed to produce cells of the SGFS. At E10, BrdU-labelled cells still resided in the VZ (Figure 12C) indicating that, in pigeon, like in chicken (LaVail and Cowan, 1971b), cells of the SGFS migrated according to an ‘inside- out developmental gradient. The fact that BrdU-labelled cells were uniformly distributed along the rostro-caudal axis at E12 (Figure 14A), suggests a temporal coincidence of the production of SGFS neurons all over the OT. The absence of identifiable gradients of cell genesis comparable to that observed in chicken (LaVail and Cowan, 1971b) is reminiscent of the much less marked central-to-peripheral gradient in the emergence of RGCs in pigeon than in chicken retina (Rodrigues et al., 2016). Taken together, our data show that the proliferation of a late pool of progenitor cells and the radial expansion of ontogenetic columns between E7 and E9 (Figure 9) have the effect of increasing the number and density of SGFS neurons who, for the most part, could be involved in retino-tectal connections. Although RGCs are the first neurons born in the retina and SGFS neurons are the last-born, genesis of the two cell types peaks around the same time – i.e., E6 in chick, E9 in pigeon (Figure 14D). This coincidence is not contingent upon axon connection that occurs at later stages of development (Rodrigues et al., 2016).

**FIGURE 14.**
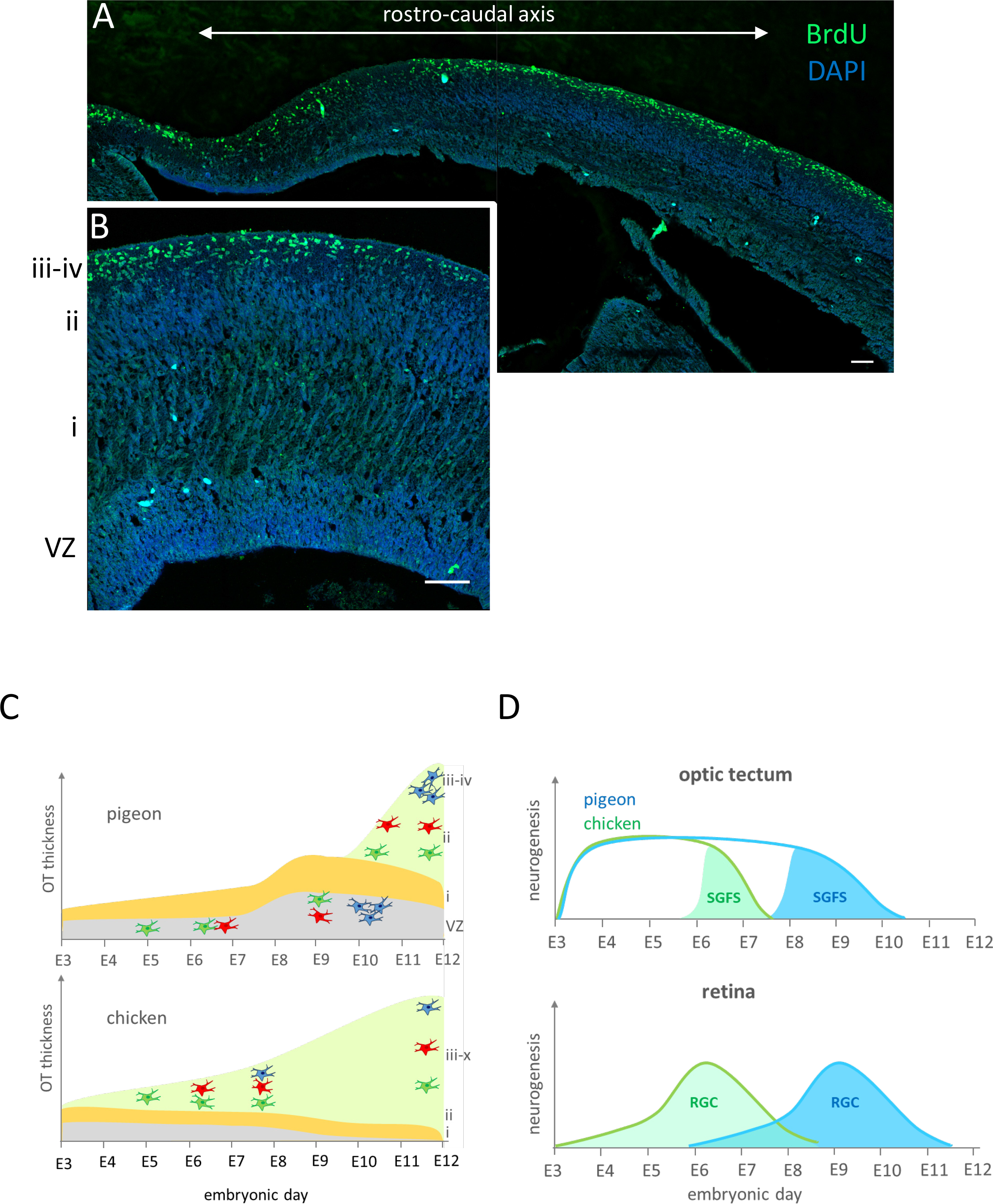
Fate and positioning of cells produced by the late pool of progenitors in pigeon. (A, B) BrdU was injected in ovo at E8. The OT was isolated at E12. A section along the rostro caudal axis (A) or a cross section (B) were processed for immunohistochemistry and counterstained with DAPI. All labelled cells are localized in the most superficial layers of the SGFS (layers iii and iv). (C) A diagram that outlines OT development in pigeon and chicken. In chicken, the newborn neuronal precursors leave the VZ, migrate through the glial fibers (layer i) to occupy their position in the outer layers (layers ii-x) according to an ‘inside-out’ developmental gradient, meaning that later-generated neurons (e.g., “blue” neuron) settle in progressively more superficial tectal layers. In pigeon, neurons do not leave the VZ until SGFS neurons arising from a late pool of progenitors have been generated. The ∼3-fold increase of neuron density in the pigeon SGFS compared to chicken results from: (1) postponed cell migration, (2) prolonged cell proliferation, (3) hindered lateral expansion of the OT. (D) A diagram that outlines neurogenesis in the OT and retina. SGFS neurons are the last-born in the OT, whereas RGCs are the first born in the retina. However, their genesis coincides in each of the two species and it is delayed by ∼3 days in pigeon. Scale bars: 50 µm (A, B).

## DISCUSSION

Our cross-species comparisons revealed that cell density in the SGFS of the OT was two to four times higher in birds with one or two foveae compared to birds with no fovea. Although this increase of cell density may look quite modest, we show that it requires spectacular changes in tissue morphogenesis and gene expression during embryogenesis. We can reasonably assume that increased densities of RGCs in the retina and of SGFS neurons in the OT of Neoaves with one or two foveae contribute to enhance their spatial and temporal visual acuity. A popular belief is that bird’s eyes send a much more processed image to the brain than our eyes send to our brain, which imply that there is less brain processing in birds than in primates (e.g., Schwab et al., 2012). However, our results are somewhat at odds with this idea as they would rather suggest that foveal vision does require an increase in the processing capacity of the brain. The emergence of the fovea in modern birds can be seen as an evolutionary breakthrough that had a decisive influence on flight, feeding habits and lifestyle. Birds became lightweight by reducing their size and evolving thin and hollow bones. The ease with which they fly and catch prey through cluttered airspaces would not have been possible without this major adaptation of their visual system. Setting up structures such as the fovea required major developmental and behavioral changes for postponing the completion of retina and brain maturation well beyond hatching, hence requiring prolonged parental care. In a wider perspective, altriciality that has developed in Neoaves ancestors may have relaxed constraints on the duration of ontogenesis, a precondition for expansion of the brain as suggested by Charvet and Striedter (2011).

The developing tectum shares features with mammalian cerebral cortex and has been viewed for a long time as a model of corticogenesis (e.g., Gray and Sanes, 1991; Hamdi and Whitteridge, 1954; LaVail and Cowan, 1971a). Gray and Sanes (1991) provided a thorough clonal analysis highlighting how migratory pathways influence cell fate determination in the developing chicken OT. They found that *“cells descended from a single progenitor begin to migrate together, but that subgroups then diverge to form three distinct streams that follow distinct guides and differentiate into distinct sets of cell types”*. In pigeon OT, cell differentiation occurs before neurons migrate out of the VZ, a fact suggesting that cell fate determination is not contingent upon cell migration in that species. One reason that could justify a drastic shift in development strategy is the necessity for pigeons to increase neuron density to levels that could not be achieved in chickens. Owing to changes in development pattern, the pigeon can increase twice the absolute cell density in its tectal lobes, thereby reaching a total number of cells equivalent to that of the chicken (∼40x10^6^ cells), yet in a tissue volume twice smaller. Cell numbers and tissue sizes increase in similar proportions in both species up to E7. Then, each species follows its own development trajectory. In chicken, tectal cells stop to proliferate and newborn neural precursors migrate to establish the outer layers. In pigeon, neurons differentiate while they are stuck in the VZ until the expansion of a pool of late progenitors does complete. This prolonged cell proliferation leads to a thickening of the VZ along its radial axis rather than to its tangential expansion. This innovative change in tissue morphogenesis may set the stage for an increase of cell densities in the outer layers once SGFS neurons did migrate (Figure 14C). We identified the earliest divergence in tissue morphogenesis when the first neurons were moving through the outer radial fibers in chicken, whereas, in pigeon, they stayed in the VZ. This observation was corroborated with the finding that a set of genes known to be involved in the regulation of cell migration were differentially expressed in chicken but not in pigeon. The uncoupling between cell differentiation and cell migration is unprecedented and raises several questions, in particular, how the VZ in pigeon provides conditions conductive to neuron differentiation, axon growth and cell positioning.

The retinas of less than 1% of the ∼10’500 bird species have been studied so far. The general trend is, however, that densities of RGCs and cones are higher in diurnal Neoaves than in Galloanserae. Higher RGC density in pigeon in comparison with chicken results from delayed neurogenesis with respect to eye growth (Rodrigues et al., 2016). While about half of retinal cells exit the cell cycle and differentiate at early embryonic stages in chicken, cells stay in the pool of proliferating progenitors for about three more days in pigeon. This strategy has the disadvantage of postponing retina development beyond hatching. In precocial Galloanserae, the need for a functional visual system at the time of hatching requires the production of RGCs at early stages, thereby de facto limiting their number. In pigeon, high level of retinoic acid and robust mitochondrial activity contribute to maintain retinal progenitors in proliferation, thereby prolonging their uncommitted status (Brodier et al., 2021; Cherix et al., 2020; Rodrigues et al., 2016). At the end of neurogenesis, the density of newborn RGCs is about three times higher in pigeon than in chicken. In pigeon, the subsequent stretching of the retinal tissue leads to a decrease in RGC density, except for a central region identified as the incipient fovea. It appears that the prolonged cell proliferation and hindered tangential tissue growth are the basic elements at work for increasing neuron densities in the fovea and in the foveal confluence of the OT.

Despite its importance, little is known about foveal vision in birds. In the human fovea, the one cone to one RGC relationship in the so-called midget pathways is presumed to be the substrate for sharp vision. However, the human fovea is not well adapted to process rapidly changing visual signals - i.e., movement perception. This low temporal acuity is what makes us see motion rather than a sequence of individual images in movies or flip books. Behavioral experiments to estimate temporal visual acuity have shown that some categories of birds can resolve alternating light-dark cycles at up to 145 Hz, which is 50 Hz over the highest frequency shown in any other vertebrate (Bostrom et al., 2016; Potier et al., 2020). Does the need to track moving objects at high-speed explain a fundamental change in the way the fovea is built and operates in birds? The three species of fast flyers catching insects on the wing (C. House-Martin, Barn Swallow, C. Swift) have fully developed central and/or peripheral foveae and the highest neuron densities in the SGFS among the studied birds. This stands in stark contrast, however, to the fact that the cone and RGC densities in the peripheral fovea of swallows and swifts are in several instances lower than in foveate birds with no peripheral fovea. The peripheral retina of the E. Sparrowhawk displays the highest RGC density among all species, yet cell density in the SGFS is lower than in species with a peripheral fovea. This somewhat paradoxical situation could indicate that for RGCs transmitting foveal visual signals, the number of tectal neurons they connect is higher than for extra-foveal RGCs. In the hawk’s peripheral retinas, the ratios of RGCs to cones are equal or close to 1 (Figure 3), suggesting the predominance of extra-foveal circuits resembling the midget pathways of primates. On the other hand, in ten out of the twelve foveae surveyed in this study, the ratios of RGCs to cones are below 0.6, suggesting that non-midget ganglion cells that receive (indirect) input from multiple cones may form alternative circuits in the bird fovea. Presence of foveal circuits with wider range of sensitivity to rapid variations in light input than in primates (Sinha et al., 2017) might provide the fovea with both high temporal and high spatial acuity, a decisive factor for birds to perceive motion in high resolution.

## ACKNOWLEDGMENTS

We are thankful to Marie Frossard for her help in tissue processing, to Philipp Delaunay and Francois Jardin for the supply of pigeon eggs, and adult pigeons (https://www.pigeonneau-normand.com/) and to Marc Plancherel for the supply of adult quails (https://www.oeuf-de-caille.com/). We wish to thank Nicolás Lendínez Bolaños (Geneva veterinary service) for his help in getting access to chicken brain tissues. Tissue processing and confocal microscopy were performed at the Bioimaging Platform of the Faculty of Sciences (http://bioimaging.unige.ch). We wish to thank Christof Bauer for his advice. RT-qPCR and RNA-Seq experiments were performed at the iGE3 genomics platform of the University of Geneva (http://www.ige3.unige.ch/genomics-platform.php). We thank Frederic Schütz (the Swiss Institute of Bioinformatics) for having made possible the completion of RNA sequencing data analysis. We are grateful to Laurent Vallotton (Natural History Museum of Geneva) for the critical reading of the manuscript and for inspiring discussions.

## DATA AVAILABILITY

Data from RNA sequencing will be made available at Gene Expression Omnibus (GEO) as soon as the paper is accepted for publication.

## MATERIALS AND METHODS

### Animals

Eyes and brains were collected from wild birds with unrecoverable injuries who have been euthanized at the rehabilitation center La Vaux-Lierre (http://www.vaux-lierre.ch). Fertilized pigeon eggs were supplied by Philippe Delaunay and François Jardin (https://www.pigeonneau-normand.com/). Chicken eggs were supplied by the animal resources centre of the University of Geneva (http://www.medecine.unige.ch/zootechnie). Adult domestic and homing pigeons were provided by François Jardin (https://www.pigeonneau-normand.com/). Adult quail and adult chicken were provided, respectively, by Marc Plancherel (https://www.oeuf-de-caille.com/) and the Fournier company (http://www.etsfournier.ch/). Birds were euthanized and tissues were collected in accordance with Federal Swiss veterinary regulations.

### Standardized tissue isolation and processing

There is a rapid decay of retinal and brain tissues in euthanized adult birds. The eyes and the brain were dissected and collected in DPBS (ThermoFisher) less than 20 minutes after death. The retinas and the OT were isolated and prefixed in 3.7% formaldehyde, 1% glutaraldehyde, 0.1 M Sodium phosphate monobasic, 0.07 M NaOH in distilled water at 4°C for 30 minutes. This 1^st^ fixation increased tissue stiffness and helped for subsequent and precise cuts. Tissue cubes of 2 to 4 millimeters side encompassing the different sectors of the retinas and OT were incubated at 4°C for a 2^nd^ period of fixation that lasted until embedding in the epoxy resin. The tissue samples were washed with 0.1 M ice cold sodium cacodylate before incubation 10 minutes in 0.8% K3Fe(CN)6 in 0.1 M sodium cacodylate and fixation in 1% osmiumtetroxide, 0.8% K3Fe(CN)6 in 0.1 M sodium cacodylate pH 7.4 for 90 minutes. They were washed with sodium cacodylate and water, stained 2 hours with 1% uranyl acetate, dehydrated in ethanol baths and incubated in propylene oxide 100% for 45 minutes, followed by propylene oxide/epon 1:1 incubation overnight. Tissue cubes were embedded in the proper orientation in epoxy resin consisting of 48% Agar 100, 18% DDSA, 31% MNA and 3% BDMA (agar scientific). Semi-thin (1 µm thick) sections were produced using a Leica UCT ultra-microtome and were mounted on glass slides (SuperFrost/plus, Assistant).

### Morphometric measurements

#### Computing cell number and cell area on tissue sections

Plastic sections 1 µm thick from embryonic and adult OT and adult retina mounted on glass slide were stained by incubation 1-2 minutes in 1% toluidine blue (Sigma Aldrich), 1% sodium borate in distilled water. Coverslips were sealed with xylene-free EUKITT® neo (O. Kindler ORSAtec). Images were acquired with a Zeiss LSM 800 microscope. With the help of tile regions, we acquired areas with dimensions that exceeded the size of an individual image field. Tiles were arranged in the form of a grid with a Zeiss image-processing software. To compute the cell number in embryonic OT and adult retina, each cell body was manually annotated by surrounding it with a closed line using an image analysis software (ImageJ/Fiji). Cells were counted in vertical frames partitioning the whole tectal or retinal section, each one of them having a width of 25-50 µm. To compute the areas of cell bodies in the retina GCL, the pixels inside the closed curve were counted using a geometry processing tool (Python library Shapely). The number of pixels was then converted to the area in µm^2^. After a test phase, we found that the counting of cell nuclei in adult OTs was more reliable when it was conducted under the microscope. The 36 resin blocks of tectal tissues (12 species and 3 sectors per species) were cut in a plane perpendicular to the tissue layers at a distance from the surface of the block that varied between 200 and 300 µm depending on the penetration depth of the fixative solution and the preservation of the tissue. Sets of four 1 µm thick sections were collected every ∼50 µm, mounted onto a glass slide (SuperFrost/plus, ThermoScientific), stained with toluidine blue, washed and air dried. From a total of 16 to 24 sections per block, 8 sections of the highest histological quality were selected. Nuclei with visible nucleoli were counted using a Nikon Diaphot inverted microscope equipped with a Nikon Fluor 40/1.30 oil objective and an eyepiece reticle (50 µm x 60 µm at the selected magnification). The counting frame was displaced across the whole area of the section encompassing the SGFS. The scanned areas varied from sectors to sectors and from specimens to specimens. On a total of 8 sections, nuclei were counted in 24 to 211 frames (referenced on the Figure 1B, x-axis). The counting of 4575 frames was carried out over a period of several months. To minimize bias in the counting procedure, tectal nuclei were counted and sometimes recounted in different specimens and sectors according to a mixed sample list. For instance, the nuclei on one section of the MT1 sector of the species x and the nuclei on one section of the MT2 sector of the species y were counted the same day, while the nuclei on another section of the MT1 sector of the species x and the nuclei on one section of the MT3 sector of the species z were counted another day.

### Isotropic suspensions of nuclei

One optic tectum (the 2 lobes) was dissected and washed in ice-cold PBS. A Dounce glass homogenizer was manually operated for shearing cells in 100 µl of lysis buffer consisting of 10 mM Tris-Cl, pH 7.5, 10 mM NaCl, 3 mM MgCl2 and 0.5% Triton X-100. Only perfectly homogenous nucleus suspensions were selected for counting. Nuclei were counted with a Neubauer chamber by phase-contrast microscopy.

### 3D reconstruction of the OT

The OT was dissected, fixed in 4% paraformaldehyde and paraffin-embedded. Properly oriented OT was cut in a plane perpendicular to the rostro-caudal axis. Every section of the ribbon of serial 10 µm thick sections was mounted on glass slides. Sections were deparaffinized and stained. Low-magnification picture of all sections were taken. To compute the surface area, each section was manually annotated by surrounding it with a closed line using an image analysis software (ImageJ/Fiji). The total volume of a dehydrated optic tectum (µm^3^) was equal to the sum of the volume of each section (i.e., surface area [µm^2^] x 10 µm).

### Reporter plasmid

The Hes5.3-GFP plasmid derive from pEGFP-C1 plasmid (Clontech). The fragment bounded by the AseI and NheI restriction sites was deleted from pEGFP-C1 and the upstream sequences of Hes5.3 (1960 bp in length, Chiodini et al. (2013)) were subcloned in the proper orientation at appropriate sites in the vector.

### In Ovo Electroporation

One hour prior to the electroporation, the eggs were rotated for half a turn so that the embryos that were sitting on the top were now at the bottom of the egg. Egg candling with a smartphone’s flashlight 1 hour later allowed to select eggs with the embryo back on the top. This test indicated that the membranes were not sticking to the shell. A piece of transparent tape was placed at the embryo position and a small hole was pierced with a forceps at the position of the air chamber. This allowed the embryo to sink before cutting the shell. A solution of DNA plasmid (1 µg/µl) and fast green dye (0.025%) in PBS was injected in the ventricle of the OT with a glass capillary mounted on a Hamilton syringe. The tweezertrodes electrodes (BTX) connected to a BTX generator were positioned on both sides of the embryonic head. We most often use the following settings: five 12.5 V/cm pulses of 50 milliseconds duration with 1 second pause between each. The window was closed using tape and the egg was incubated until reaching the desired stage.

### Confocal imaging of the optic tectum

Electroporated OTs were fixed with 4% paraformaldehyde for 20 minutes and were mounted on concave glass slides. Coverslips were sealed after addition of DABCO with 50% glycerol (Sigma-Aldrich) in DPBS. Images were acquired with a Zeiss AXIO Imager Z1m microscope.

### BrdU labelling

#### In ovo

Pigeon embryos received a single dose of 5-Bromo-2′-deoxy-Uridine (100 µl of PBS containing 10 mM BrdU) into the chorioallantois, through a window made in the shell the date of injection. The window was sealed with transparent tape, and the egg returned to the incubator.

#### Ex vivo

The optic tectum explants were incubated in 35 µM BrdU, in Dulbecco’s Modified Eagle Medium (DMEM), 10% fetal bovine serum (FBS) for 1 or 19 hours. The explants incubated for 1 hour in BrdU were transferred in DMEM, 10% FBS for 1 hour, rinsed in PBS, fixed in 4% paraformaldehyde and embedded in paraffin. Tectal cells from explants incubated for 19 hours in BrdU were dissociated in 0.1% trypsin in Hanks’ Balanced Salt Solution (HBSS) without Ca^++^ and Mg^++^. 50 µl of DNase and 500 µl of FBS were added 5 and 15 minutes later. The cell suspension was centrifuged and the pellet was resuspended in DMEM. Cells were counted with a Neubauer chamber and 10^5^ cells in a 100 µl drop were deposited on a polyornithine-coated plastic chamber slide (Lab-Tek). Slides were incubated at 37°C for 1 hour to allow cell attachment, fixed in 4% paraformaldehyde for 20 minutes, rinsed in PBS and air dried.

#### Immunostaining

Deparaffined sections (8 µm thick) or dissociated cells were rehydrated, rinsed in PBS and incubated 10 minutes (dissociated cells) or 15 minutes (tissue sections) in 4N HCl. After neutralization in PBS, 0.5% BSA, 0.1% Triton X-100, cells and sections were processed for BrdU immunodetection following the manufacturer’s recommendations (5-Bromo-2′-deoxy- uridine Labelling and Detection Kit I, Roche). The slides were mounted for fluorescent microscopy in ProLong® Diamond Antifade Mountant with DAPI (Life Technologies).

### RT-qPCR

RNA from whole OT was extracted using TRIzol reagent (ThermoFisher) according to product manual in triplicate. Primers were designed using NCBI primer blast and Primer 3 websites. Primers were ordered from Microsynth. Primers were tested using RNA from total retina extracted with TRIzol. RNA quantification was done with spectrophotometer and Qubit 2.0 (ThermoFisher) and RNA quality was checked using BioAnalyzer 2100 (Agilent). RT was performed using Takara PrimeScript reverse transcriptase prior to qPCR. In samples with delta-Ct values > 0.5 across the three technical replicates, the most extreme value was removed. Quantities were calculated as 2^(Ctmin-CT). The Average-Quantity was calculated for each sample. GeNorm (Vandesompele et al., 2002) was used to determine the best normalization genes. A normalization factor was calculated as Normalization-Factor ¼ Geometric Mean [Average-Quantity of Normalization Genes] and was used to calculate the normalized quantity: Normalized-Quantity ¼ Average-Quantity/Normalization-Factor. The fold change was calculated as: Fold-Change ¼ Mean [Experiment condition]/Mean [Baseline condition]. P-values were calculated using a Welch t-test to compare baseline and experiment samples (in log 2 scale). RT-qPCR were performed using the primers listed in Table S3. All specific primers for chicken and pigeon listed in Table S3 were tested for efficiency.

### RNA-Seq

#### Tissue collection

We collected 5 biological replicates for each of the three developmental stages (E6, E8 and E10) of pigeon and chicken OT (30 samples). Total RNA was extracted and quantified with a Qubit (fluorimeter from Life Technologies) and RNA integrity assessed with a Bioanalyzer (Agilent Technologies).

#### RNA sequencing

A total of 30 RNA sequencing libraries was generated. Libraries were prepared with TruSeq Stranded RNA Library Prep Kit (Illumina) and sequenced on multiple lanes of an Illumina HiSeq2500 platform, generating paired-end 100-cycle reads for each sample. The quality of sequence was assessed by FastQC (version v0.11.4, available online at: http://www.bioinformatics.babraham.ac.uk/projects/fastqc/ (Andrews, 2010). Purity-filtered reads were adapters- and quality- trimmed with Cutadapt (v1.8) (Martin, 2011). Reads matching to ribosomal RNA sequences were removed with fastq_screen (v. 0.9.3). Remaining reads were further filtered for low complexity with reaper (v. 15-065) (Davis et al., 2013).

#### Chicken gene annotation

Gallus gallus reads were then aligned against Gallus_gallus-5.0.92 genome release using STAR (v.2.5.2b) (Dobin et al., 2013). The number of read counts per gene locus was summarized with htseq-count (v0.6.1) (Anders et al., 2015). Quality of the RNA-seq data alignment was assessed using RSeQC (v. 2.3.7) (Wang et al., 2012).

#### Pigeon gene annotation

Pigeon genome (Cliv_1.0 assembly from 2014-02-18) was obtained from the (GIGA)n database (gigadb.org). Since the pigeon is not a model organism, we annotated the genes by comparing Columba livia gene sequences to Gallus gallus genes. We realized a sequence mapping between pigeon and chicken using blast by keeping the maximum score hit.

#### Differential Expression Analysis

To assess differential expression between embryonic development stages of the pigeon and chicken, we compared the foldchange of significantly differentially expressed genes between E6 and E8 and between E8 and E10. Data analysis has been performed with the R Bioconductor package DESeq2 (version 1.14.1) (Love et al., 2014). Differentially expressed genes (DEGs) were identified at the Benjamini-Hochberg (BH) adjusted P < 0.05 level, using Wald test.

### Statistical analysis of morphometric data

All computations and graphics were done using Python v3.9.2 (libraries: Matplotlib v3.4.2., Numpy v1.20.2, Scipy v1.6.3, Shapely v1.8a3). The Kernel Density Estimation (KDE) used for drawing the violin plots (Figures 1B, 3B) consists in extracting an empirical probability density function from the data (i.e., a non-negative function whose area under the curve equals one). Given a set of *n* data points *x*_1_, …, *x*_*n*_, the density estimate at a point *x* is obtained as

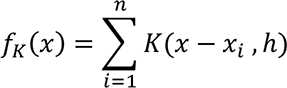

where *K*(*x*, ℎ) is a positive function controlled by a bandwidth parameter ℎ > 0. One can think of the kernel *K*(*x*, ℎ) as a smoothing sliding window of width ℎ which can have different shapes. The shape of the kernel reflects the weight of a data point *x*_*i*_ located at a given distance of the estimated point *x*. We used a kernel with a gaussian shape :

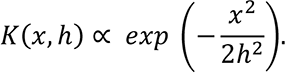

The kernel is appropriately normalized so that the area under the curve *f*_*K*_ equals one. The bandwidth parameter ℎ was set according to Scott’s Rule :

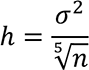

where σ is the computed standard deviation of the data.

The red curve *g* (Figure 5) is obtained by smoothing the binned values *x*_1_, …, *x*_*n*_ with a gaussian kernel

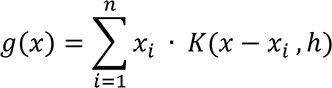

where *K*(*x*, ℎ) is the same gaussian kernel than for the KDE estimation with a bandwidth empirically set to ℎ = 5.

## SUPPLEMENTARY DATA

**TABLE S1.** List of specimens used in the study. Abbreviations: nd, not determined; F, female; M, male; J, juvenile; A, adult; Tel, telencephalon; OT, optic tectum; diam, diameter.

| Species | Species (EN) | Age | Sex | Body weight (g) | Brain weight (g) | Eye diam. (mm) | Tel. length (mm) | Tel. width (mm) | Tel. height (mm) | OT length (mm) | OT width (mm) | OT height (mm) |
| --- | --- | --- | --- | --- | --- | --- | --- | --- | --- | --- | --- | --- |
| *Delichon urbicum* | Common house martin | J | nd | 17,33 | 0,49 | 9 | 9 | 7 | 5 | 4 | 2 | 3 |
| *Hirundo rustica* | Barn swallow | J | M | 15,06 | 0,64 | 9 | nd | nd | nd | 5 | 2,5 | 4 |
| *Falco tinnunculus* | Common kestrel | A | M | 158,53 | 3,8 | 20 | 16 | 13 | 10 | 9 | 5 | 7 |
| *Apus apus* | Common swift | A | F | 32 | 0,86 | 12 | 10 | 8 | 6 | 5 | 3 | 4 |
| *Columba livia* | Pigeon | A | nd | 665 | 2,76 | 15,8 | 13,7 | 9,5 | 8,7 | 8,5 | 5,8 | 6,1 |
| *Matocilla cinerea* | Grey wagtail | J | nd | 15,73 | 0,65 | 8 | nd | nd | nd | 5,5 | 3 | 4 |
| *Accipiter gentilis* | Northern goshawk | J | M | 776 | 7,96 | 27 | 21 | 18 | 13 | 12 | 7 | 9 |
| *Accipiter nisus* | Eurasian sparrowhawk | A | M | nd | nd | 19 | 15 | 12 | 8 | 9,5 | 5,5 | 7 |
| *Milvus migrans* | Black kite | A | M | 807 | 7,51 | 24 | 19 | 14 | 12 | 10 | 6 | 9 |
| *Strix aluco* | Tawny owl | A | nd | 479 | 9 | 29 | 23 | 18 | 15,5 | 10,5 | 5 | 8,5 |
| *Gallus gallus* | Chicken | A | nd | nd | 3,53 | 21 | 18 | 11 | 10 | 10 | 5 | 7,5 |
| *Coturnix japonica* | Japanese quail | A | F | 324 | 0,86 | 11 | 10 | 7 | 6 | 6,5 | 4 | 5 |

**TABLE S2.** Cell counts per OT (2 lobes) obtained with isotropic suspensions of cell nuclei. Each value is the mean ± s.d. of biological triplicates.

|  |  | E3 | E4 | E5 | E6 | E7 | E8 | E9 | E10 | E11 | E12 | E13 | E14 | E15 |
| --- | --- | --- | --- | --- | --- | --- | --- | --- | --- | --- | --- | --- | --- | --- |
| Chicken | cell number | 843 667 | 2 643 333 | 8 313 333 | 26 495 000 | 39 061 250 | 44 398 333 | 42 945 000 | 47 667 500 | 45 539 000 | 45 809 000 | 45 236 667 | 44 003 333 | 45 300 000 |
|  | stdev | 240666,44 | 684855,22 | 938580,49 | 3599161,9 | 4536254,7 | 2505944,6 | 2966315,1 | 265835,95 | 2124474,8 | 483996,9 | 2973589,3 | 1231273,1 | 2219797,3 |
|  | divisions in 24h | n/a | 1,6 | 1,7 | 1,7 | 0,6 | 0,2 | 0 | 0,2 | 0 | 0 | 0 | 0 | 0 |
| Pigeon | cell number | 485 140 | 888 133 | 1 979 250 | 7 173 333 | 17 790 000 | 26 900 000 | 34 348 333 | 35 620 000 | 40 593 333 | 39 698 333 | 41 680 000 | 47 423 333 | 44 023 125 |
|  | stdev | 115390,2 | 92606,983 | 391478,71 | 2904140,7 | 2721635,5 | 1317725,3 | 1551856,4 | 3267292,5 | 3371977,7 | 3696729,4 | 3974295 | 1875535,5 | 3452338,2 |
|  | divisions in 24h | n/a | 0,9 | 1,2 | 1,9 | 1,3 | 0,6 | 0,4 | 0,1 | 0,2 | 0 | 0,1 | 0,2 | 0 |

**TABLE S3.**
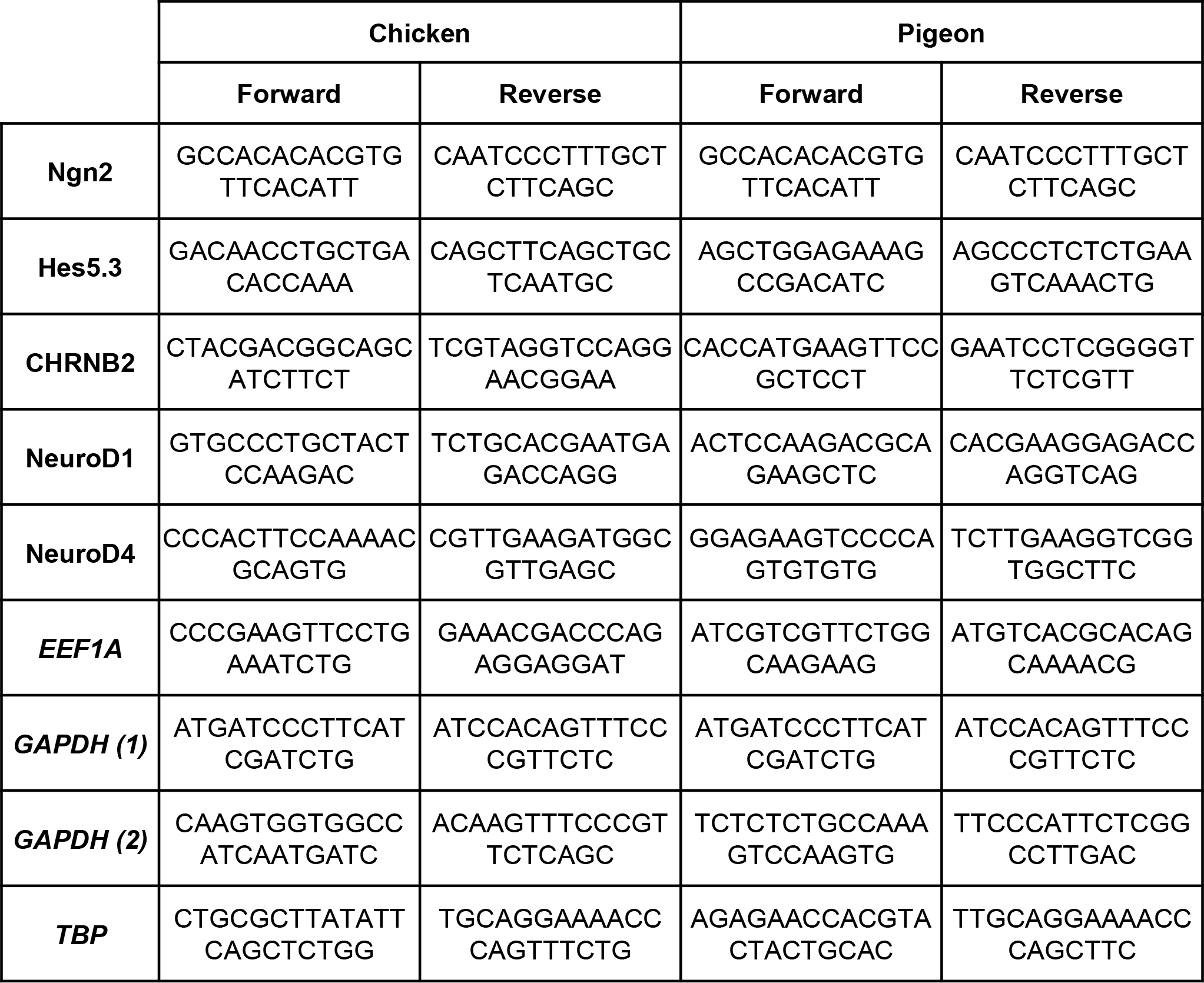
List of primers used in this study for RT-qPCR (transcripts). Genes used for normalization of RT-qPCR are in italic.

